# Autonomic Indicators of Self-Transcendence: Insights from the Numadelic VR Paradigm

**DOI:** 10.1101/2025.11.11.686736

**Authors:** Valerie Bonnelle, Giulia Parola, Catherine Andreu, Joseph L Hardy, Justin Wall, Christopher Timmermann, Ausiàs Cebolla, Maja Wrzesien, David R Glowacki

## Abstract

Self-transcendent experiences (STEs) offer profound and beneficial shifts in perspective, yet remain largely inaccessible outside elite contemplative or pharmacological contexts. Neural measures have deepened our understanding of these states, but their cost and limited ecological validity constrain broader application. This study evaluates heart rate variability (HRV) amplitude, a measure reflecting dynamic sympathovagal engagement, as a cost-effective and sustainable physiological marker of STE during ‘numadelic’ virtual reality (VR) experiences. The unique numadelic aesthetic combined with multi-person can induce STE through dissolving self-boundaries and fostering embodied presence.

Building on previous work linking non-ordinary states of consciousness (NOSC) with autonomic nervous system activity during psychedelic drug administration, we tested the hypothesis that HRV amplitude may reflect STE depth and relate to affective and relational outcomes during non-drug numadelic VR experiences. Specifically, ninety-six participants engaged in guided meditation within either a numadelic VR setting or a non-VR audio-guided group format. Physiological data (cardiac activity and respiration) were recorded during the meditation, alongside psychological assessments pre- and post-session.

Findings confirm that HRV amplitude measured during numadelic VR correlates with subjective STE ratings. It also relates to compassion traits, and emotional improvement following VR practice. Additional analysis of data from a prior psychedelic drug study further validated the relevance of this measure across methods of inducing NOSCs.

These results advance the psychophysiological mapping of STEs and highlight HRV amplitude as a potential real-time biomarker which may help to guide participants toward self-transcendent states within adaptive environments. By integrating contemplative science with immersive design, this work supports the development of scalable tools that enhance both access to and understanding of STEs.

## Introduction

Experiences like witnessing breathtaking natural phenomena—standing before a vast mountain range or under a star-filled sky—being profoundly moved by music, art, or literature, or feeling deep love and care for others, are moments when our usual sense of self transcends its boundaries. Such experiences often lead to transformative shifts in perspective, broadening our understanding of our place in the world, while fostering hope, peace, unity, and enhanced compassion (Koltko-Rivera, 2006; Stellar et al., 2017).

According to Reed, self-transcendence can be defined as an “*expansion of self-conceptual boundaries multidimensionally: inwardly (e.g., through introspective experiences), outwardly (e.g., by reaching out to others), and temporally (whereby past and future are integrated into the present)”*, that also includes a transpersonal expansion component, which involves connecting to what is experienced as a higher power or a larger reality (Reed, 1991, p 64). Self-transcendent experiences (STEs) have been a cornerstone of human history, deeply embedded in philosophical, spiritual, and psychological traditions (Wong, 2016). They constitute central element of traditional practices such as meditation, rituals, and communal ceremonies sought to dissolve ego boundaries, connecting individuals to a higher purpose or collective identity (Frey and Vogler, 2018). In contemporary Western contexts, however, such practices have become less accessible and often marginalised, possibly contributing to a growing sense of disconnection and existential distress (Turkle, 2011; Osler, 2022).

Amid global challenges and the erosion of communal bonds, there is renewed interest in contemplative modalities and non-ordinary states of consciousness (NOSC) as potential pathways to meaning, resilience, prosociality, and psychological wellbeing (Franco Corso et al., 2023; Grof and Grof, 2023; Ko et al., 2022). The psychedelic ‘renaissance’ that has unfolded in the past decade, alongside the emergence of mindfulness and compassion based programs, reflect this trend toward a framework that re-introduces STEs in our approach to medical care and personal growth (Hadar et al., 2023). However, the accessibility and broader adoption of these practices remain limited. The democratisation of psychedelics requires careful navigation due to the challenges associated with ensuring their safe, ethical, and supervised use (Pilecki et al., 2021). As for meditation practices, while increasingly popular, they often require sustained dedication to cultivate the depth and maturity necessary for the emergence of STEs.

The present study explores novel tools for eliciting and monitoring STEs through the integration of immersive virtual reality (VR) and physiological monitoring. VR enables the creation of immersive, interactive environments that can profoundly reshape users’ perception, embodied presence and agency (i.e. the user’s felt ability to act, influence, and make meaningful choices in VR) (Slater and Sanchez-Vives, 2016; Suzuki et al., 2023). These altered experiential states have been shown to facilitate STEs, offering pathways to expanded awareness and emotional insight (Quesnel and Riecke, 2018; Vidal et al., 2024). As such, VR holds promise as a catalyst for personal transformation with potential for therapeutic applications (Riva et al., 2016). VR offers the tantalizing possibility for constructing immersive sensory environments that can represent a vast range of phenomena, limited by little more than imagination. Nevertheless, the vast majority of VR content utilized in research contexts is relatively limited – i.e., it tends to follow the dominant aesthetics of the metaverse and maintain fidelity to the sorts of visual scenes which people can encounter during their day-to-day experience (Glowacki, 2024). Among existing VR experiences, those designed within the ‘numadelic’ aesthetic take a dramatically different design approach, representing human bodies as unbounded essences composed of light and energy (NB: The term *numadelic* is derived from Greek words ‘pneuma’, meaning ‘breath’ or ‘spirit’, and ‘Delein’, meaning ‘to revel’ or ‘to manifest’). Within VR, the numadelic aesthetic, combined with a platform supporting multi-users experiences, has been shown to elicit STEs of comparable phenomenological intensity to those induced by moderate doses of psychedelics (Glowacki et al., 2022, 2020). In multi-person numadelic VR environments, participants experience themselves as luminous, energetically diffuse beings. This blurring of self/nonself distinctions can foster a sense of interconnectedness and self-transcendence (Glowacki, 2024; Vidal et al., 2024).

Maximizing the depth of STEs during a numadelic journey depends on optimizing embodiment and presence, which heavily relies on emotional engagement in the experience (Riva et al., 2016; Yaden et al., 2017). VR development often employs a user-centred design approach, but this reliance on subjective user accounts can result in limited feedback accuracy and poor temporal resolution (Napa Scollon et al., 2009; Vlahovic et al., 2022; Winkler and Appel, 2024), as well as demand characteristics and behavioural compliance, that is the tendency for participants to be influenced by what they think experimenters are looking for (Suzuki et al., 2023). The use of physiological measures as proxies of participants’ subjective experience appears as a promising approach to collect useful information about participants’ emotional state during their experience, for instance offering a real-time, objective way to monitor STEs as they unfold. In addition, such measures may ultimately be integrated as a real-time biofeedback which modifies the VR environment, helping guide participants toward optimal states that may be conducive to STEs.

Can STEs be reflected in physiological state? The idea that mental states are mirrored in the body is a core concept in the James-Lange theory of emotion, which posits that emotions are the result of interpreting our bodily reactions to stimuli (James, 1994). While the original formulation may be overly simplistic, contemporary perspectives in embodied cognition and affective neuroscience emphasize emotions as embodied phenomena that regulate adaptive behavior and connect mental processes to sensory and motor experience (Laird, 2007; Laird and Lacasse, 2014; Oosterwijk et al., 2012). Motivational and appraisal models further frame emotions as embodied states that prepare the organism for adaptive action (Scarantino, 2014). The autonomic nervous system (ANS) provides the primary physiological substrate for these embodied states, orchestrating cardiovascular, respiratory, and visceral adjustments. These bodily changes can become subjectively accessible through interoceptive awareness (Cameron, 2001). Regulation of this brain-body loop is coordinated by the central autonomic network, a distributed network including the insula, anterior cingulate, amygdala, and prefrontal cortex, that integrates autonomic activity with emotional and cognitive processes. Within this framework, the vagus nerve plays a pivotal role in transmitting visceral information (Porges, 2009), while interoceptive pathways provide the substrate for body-mind integration (Craig, 2002; Critchley, 2005). The ANS thus emerges as a dynamic mediator between emotion generation and regulation, linking bodily states with cognition, decision-making, and self-awareness.

The ANS consists of two branches: the sympathetic (SNS) and parasympathetic (PNS) nervous systems, which are involved in responding to and regulating internal and external stimuli. The sympathetic branch activates the ‘fight or flight’ response, releasing adrenaline and noradrenaline to increase arousal by elevating heart rate (HR), blood pressure, and respiration rate (Jänig, 2003). These changes prime the body for heightened sensory awareness and physical readiness for action by directing energy to muscles, lungs, and bottom-up attentional processing brain centres, while reducing activity in less urgent systems like digestion and default mode network (Beissner et al., 2013). This high arousal state is also a crucial component of emotional processing, ensuring an adaptative response, particularly in situations involving intense emotions such as stress, fear, or excitement, where the body mobilizes resources to address perceived challenges, threats, or opportunities.

The parasympathetic branch, on the other hand, facilitates the ‘rest and digest’ response by releasing acetylcholine, which lowers HR, decreases blood pressure, and promotes relaxation (Jänig, 2003). It supports the conservation of energy, digestion, and the restoration of bodily tissues. This branch plays a key role in calming or regulating emotional states, helping the body recover from arousal caused by the sympathetic system, and fostering feelings of safety and wellbeing that form the basis of healthy humans’ interactions (Duarte and Pinto-Gouveia, 2017; McCorry, 2007; Pinna and Edwards, 2020; Porges, 2022).

While it has been assumed in the past that the two branches of the ANS act solely in a reciprocal manner, with increases in sympathetic activity paired to decreases in PNS activity (Cannon, 1914, 1915; Fulton, 1949), more contemporary research indicates that activity in both branches can vary independently, defining a two-dimensional regulatory space rather than a single axis of reciprocal opposition between the two branches (Berntson et al., 1993). Both branches are integrally involved in emotional regulation and processing, and their dynamic interplay is essential not only for maintaining homeostasis but also for adapting to the emotional and social demands of daily life (Kreibig, 2010; Siegel et al., 2018; Stifter et al., 2011).

As well as being involved in emotional processing, the state of the ANS has also been found to affect and to be affected by contemplative practices. A number of meditation practices have long been recognized for their relaxing effects (Davidson, 1976; Kushner and Marnocha, 2008), which are largely mediated by the activation of the PNS, providing physiological restoration, or homeostasis (Benson et al., 1974). Noteworthily, certain forms of meditation, particularly those associated with enhanced affective and prosocial processing, such as compassion-based practices (Dahl et al., 2015), do not solely induce relaxation. Instead, these practices have been linked to a unique physiological state involving the simultaneous engagement of both SNS and PNS components (Léonard et al., 2019; Lutz et al., 2009; Sezer and Sacchet, 2025). This dynamic interplay results in a balanced and positively regulated form of arousal that promotes prosocial motivation and engagement. In addition, in traditional Buddhist frameworks, meditation is understood as a state of calm yet vigilant awareness—one that requires careful balancing to avoid the extremes of hyperarousal (manifesting as restlessness) and hypoarousal (such as drowsiness or sleep) (Britton et al., 2014). This state, described as ‘relaxed alertness’, corresponds to a unique combination of a still, relaxed body and an active, focused mind. A possibly over-simplified view would be that while early stages of meditation practice mostly promote relaxation through PNS activation, more advanced practice may involve mastering a balanced state between PNS and SNS activity (Sezer and Sacchet, 2025).

Beyond meditation, research into the physiological correlates of STEs remains in its early stages. However, several studies suggest that certain types of NOSC can be induced through hyperarousal. This phenomenon has been intuitively understood for millennia, as evidenced by the widespread use of various high arousal trance-inducing practices in ancient traditions (e.g. hyperventilation, fasting, sweat-lodge, dance, drumming) (Gosseries et al., 2024; Oswald et al., 2023; Walter and Altorfer, 2023).

How can hyperarousal lead to spiritual experiences? Bonnelle et al. (2024) have proposed that it may be related to the so-called vagal rebound that follows an intense stress response. This process involves a rapid physiological shift: during acute stress, PNS activity is withdrawn, but in the recovery phase it reactivates, often surpassing baseline levels (Mezzacappa et al., 2001). A well-documented example is HR recovery following aerobic exercise. During exertion, strong SNS activation and PNS (or vagal) withdrawal allow the body to meet its metabolic demands. Immediately afterward, PNS reactivation produces rapid HR deceleration to promote a return to homeostasis. Importantly, SNS activity can remain transiently elevated during this recovery phase, creating a brief state of co-activation between the two branches of the ANS (Perini and Veicsteinas, 2003; Weissman and Mendes, 2021). This unusual and transient physiological state of heightened, yet regulated, arousal may provide the adequate physiological conditions for STEs. Importantly, sympathovagal co-activation may only be one physiological manifestation of such state, which reflects a complex interplay of bottom-up visceral signaling and top-down central processing, temporarily reshaping mind–body integration by balancing the usual dominance of one autonomic branch over the other (Thayer et al., 2009).

Consistent with this ‘vagal rebound’ hypothesis, previous work has shown that individuals who reported the most profound spiritual and insightful experiences following the administration of the psychedelic drug N,N-Dimethyltryptamine (DMT) - which is associated with a pronounced increase in sympathetic activity immediately post-administration - demonstrated a higher state of co-activation of the SNS and PNS during their experience (Bonnelle et al., 2024). Another potential example of this state of sympathovagal interplay may be found in aesthetic chills. This psychophysiological response, often associated with the experience of awe and self-transcendence (Christov-Moore et al., 2024), seems to reflect a state of high sympathetic arousal experienced in a context of awe-induced PNS activation (Gordon et al., 2017; Jain et al., 2023). Finally, compassion – defined as the feeling that arises in witnessing another’s suffering and that motivates a subsequent desire to help – is another type of STE deeply associated with the state of the ANS (Di Bello et al., 2020; Goetz et al., 2010). While traditionally described as a vagally mediated response, contemporary models highlight the dynamic interplay of both autonomic branches (Di Bello et al., 2021). This view captures the phenomenology of compassion as a balance between activation (empathic presence and readiness for action) and regulation (managing difficult emotions to sustain connection and reduce empathic distress) (Condon and Barrett, 2013; Gilbert and Van Gordon, 2023; Strauss et al., 2016). At the autonomic level, this requires flexible engagement of both branches: rapid sympathetic activation to support empathy and mobilization, and parasympathetic activity to facilitate emotional regulation (Critchley, 2005; Di Bello et al., 2021; Laborde et al., 2017; Thayer and Lane, 2009).

This state of sympathovagal dual engagement can be measured through heart rate variability (HRV). HRV refers to the variations in time between successive heart beats, which reflects the dynamic interplay between the SNS and PNS (Rajendra Acharya et al., 2006). The Neurovisceral Integration Model suggests a top-down view where HRV reflects the prefrontal cortex’s regulation of the ANS, enabling effective emotional regulation and fostering positive emotional experiences (Thayer and Lane, 2000). This relationship, however, may be bidirectional. For instance, elevated HRV is linked to positive affect through enhanced self-regulation capacity and stress adaptivity. Yet, positive affect broadens coping mechanisms, improves stress recovery, and ultimately promotes the autonomic flexibility required for higher HRV (Schneider et al., 2025).

To capture this complex interplay, it is important to consider that the temporal dynamics of the ANS branches differ significantly. The SNS’s influence on the heart is slower acting and sustained, precluding substantial beat-to-beat changes (Jose and Collison, 1970). Consequently, high-frequency (HF) cardiac rhythms are mediated primarily by PNS innervation, while low frequency (LF) reflects a dynamic mixture of SNS and PNS contributions (Quigley et al., 2024). The former measure (HF-HRV), as well as other standard time-domain measures focusing on fast beat-to-beat fluctuations (e.g. RMSSD) are often used to measure the specific contribution of the PNS in cardiac activity (Shaffer and Ginsberg, 2017). In this study, we were interested in evaluating the overall engagement and interplay between the two ANS branches and its involvement in STEs, in a way that allows dynamically tracking and objectively measuring such experiences. For that specific purpose, we computed a new measure of sympathovagal engagement estimated from the overall envelope of HR fluctuations averaged over several heart beats (Figure 1).

**Figure 1:**
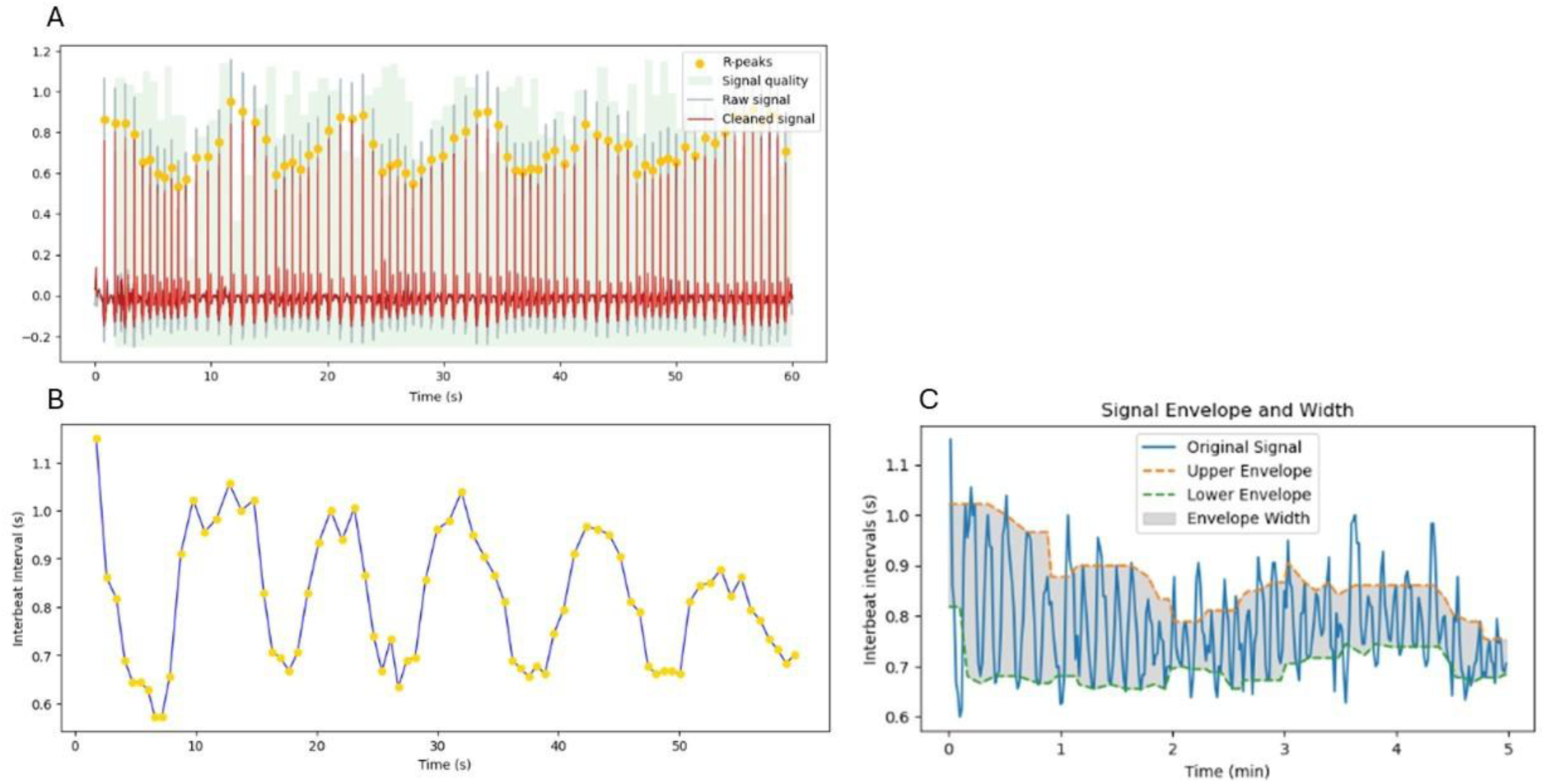
A) Example of one minute of ECG signal (red), with R-peaks identified by yellow points. B) Interbeat intervals (intervals between two consecutive R-peaks) plotted over time (NB: this is heart rate variability (HRV)). C) Example of interbeat intervals, or HRV plot, over a 5 min period. The upper and lower envelopes of the HRV plot can be derived (as described in Method) in order to estimate local HRV amplitude, a measure reflecting sympathovagal engagement.

### Study objectives

Our study pursued four specific aims. First, after confirming that numadelic VR reliably elicits deeper STEs than non-VR meditation, we tested our primary hypothesis: that HRV amplitude (a measure that captures the dynamic interplay between the two ANS branches) would reflect subjective experience of self-transcendence during the VR meditation.

Second, because STEs are often associated with lasting improvements in mood and relationality (Kitson et al., 2020; Yaden et al., 2017), we examined whether HRV amplitude measured during VR sessions correlated with post-practice changes in these dimensions, thereby testing the potential of this measure as an objective indicator of emotional and social benefits post VR experience.

Third, recognizing that compassion and self-transcendence mutually reinforce one another (Crockett and Lockwood, 2018; Kang, 2019), we examined the role of compassion traits. Our goal was to assess whether dispositional compassion was associated with STE depth and HRV amplitude during the experience, providing insight into how pre-existing traits might influence autonomic response during contemplative practices.

The final part of this study was conducted as an *a posteriori* analysis to evaluate the robustness and generalizability of our main findings. This approach acknowledges the considerable variability inherent in studies investigating the relationship between HRV and psycho-emotional states, which often reflects diverse methodologies and contexts, thus complicating the broader extrapolation of results (Schneider et al., 2025). To address this concern, we revisited data from a previously published study of DMT-induced NOSC. DMT is a short-acting serotonergic psychedelic that offers a unique opportunity to investigate shifts in autonomic activity and their relation to STEs as its short duration of action (∼15-20 min) allows drawing more accurate relationships between subjective reports of acute effects and physiological measures. This additional analysis allowed us to test whether the autonomic signature of STEs observed in numadelic VR, specifically HRV amplitude, also emerges in a pharmacologically induced context. By extending the analysis across distinct modalities, we aimed to strengthen the case for HRV amplitude as a scalable, cross-context biomarker of STE depth, potentially unifying diverse approaches to eliciting STEs.

## Methods

### Study design

The main study from which this work stems (Validation of the efficacy of Numadelic VR inducing STE, currently under preparation) was preregistered on the Open Science Framework prior to data collection (https://doi.org/10.17605/OSF.IO/YNPCK), and employed a randomized controlled design with two groups, an experimental group participating in a group numadelic VR experience as implemented within the AnumaXR app (available on the Meta Quest App Lab) on Meta Quest 3 wireless head mounted displays (HMDs), and the control group in a traditional group setting (e.g., recorded meditation), both practicing a guided meditation in sub-groups of four. Within the AnumaXR app, group VR experiences (wherein each participant occupies the same virtual space) are possible by connecting each headset to the same cloud-mounted server. The present study constitutes an exploratory analysis addressing secondary objectives outlined in the pre-registration.

Each participant underwent assessments before the practice and immediately after the practice. Data collection involved administering a battery of questionnaires at baseline and post experience, and recording physiological measures (ECG and Respiration) during the practice session, and during two rest periods before and after the session.

### Participants

This study was approved by the Ethics Committee of the University of Valencia, Spain (registration number: 2024-PSILOG-3349305). All activities conducted in studies involving human participants adhered to the principles outlined in the 1964 Helsinki Declaration and its subsequent revisions or adhered to equivalent ethical standards.

In total, 96 participants (12 sub-groups of 4 for each condition) over 18 years old and proficient in Spanish were recruited through flyers and social media. Exclusion criteria were as follows: (a) current diagnosis of a psychological disorder (e.g., depression, anxiety), (b) use of substances altering the nervous system, (c) cardiac conditions, and (d) current or past diagnosis of post-traumatic stress disorder (PTSD), psychosis and personality disorders. Upon enrolling to the study, participants were randomly assigned to one of the conditions using https://www.randomizer.org/ software.

Three participants in the VR group were excluded from further analysis; one did not complete the experimental session, one reported that they felt confused by the questionnaires after the experience, and one experienced an adverse reaction halfway through the session and significant discomfort with the headset (Final VR group: n= 45, mean age =27 ± 11 years, 28 males). Two people did not show up on their session day in the control group, and had to be replaced with research team members, whose physiological data were collected but not included in the analysis (Final Control group: n=46, mean age = 23 ± 7 years, 34 males). The control group was significantly younger (Mann-Whitney U test, p=0.046), but there was otherwise no significant difference in gender, Body-Mass Index (BMI) or meditation experience between the two groups.

### Experimental Procedure

Participants completed baseline questionnaires and attended group experimental sessions (blinded for group assignment until the beginning of the session) in sub-groups of four. The experimental sessions were carried out by a facilitator and a person in charge of physiological data collection (both not blind for participants group assignments). The practices, in both VR and control groups, included three distinct phases during which participants remained seated: preparation, meditation, and integration/discussion. Preparation involved facilitator introduction, explanation of the meditation, intention-setting, and guided breathing practices. Following this initial phase, the facilitator started a pre-recorded guided meditation lasting approximately 25 minutes, composed of seven distinct chapters described in Table 1. Last, participants were invited to share their experiences with the group. Practices were preceded and ended with five minutes of rest periods, with eyes closed in order to collect physiological data (without the VR headset for the VR group). Our experimental procedure is summarized on Figure 2.

**Figure 2:**
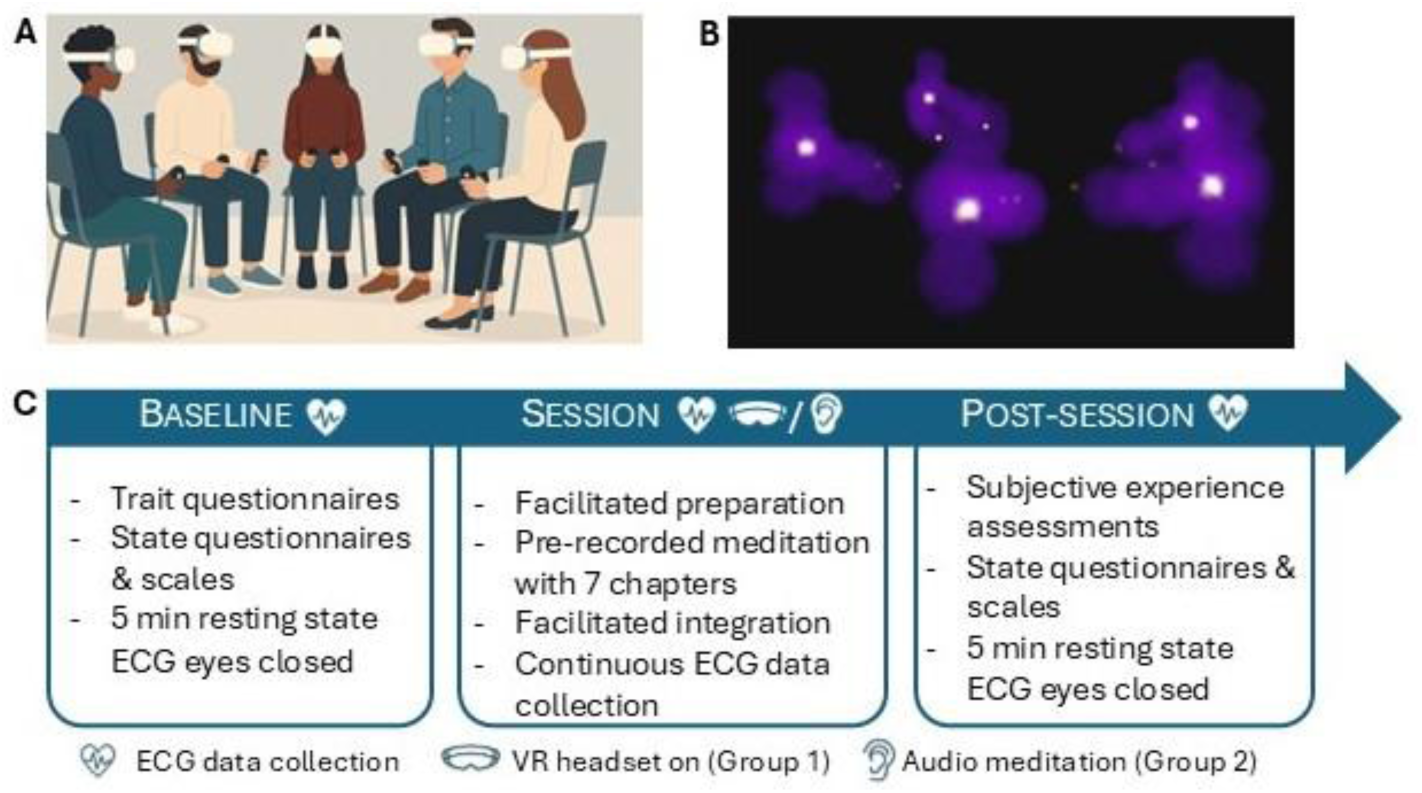
Experimental procedures. A) Illustration of participants (n=4) and facilitator during a VR session. B) Visual representation of the virtual space with 4 participants in their numadelic avatars. C) Study design on data collection days. The list of questionnaires and scales used at baseline and post-session can be found in the ‘Questionnaires’ section of the Methods. Physiological data analysis presented in this article is focused on the Pre-recorded meditation sequence described in Table 1.

**Table 1:**
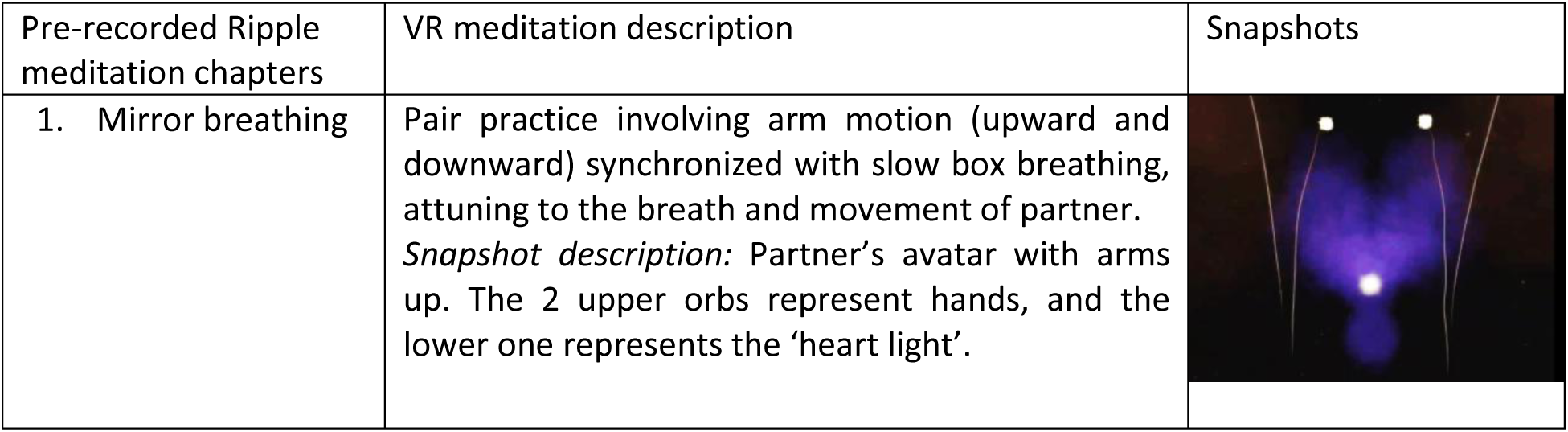
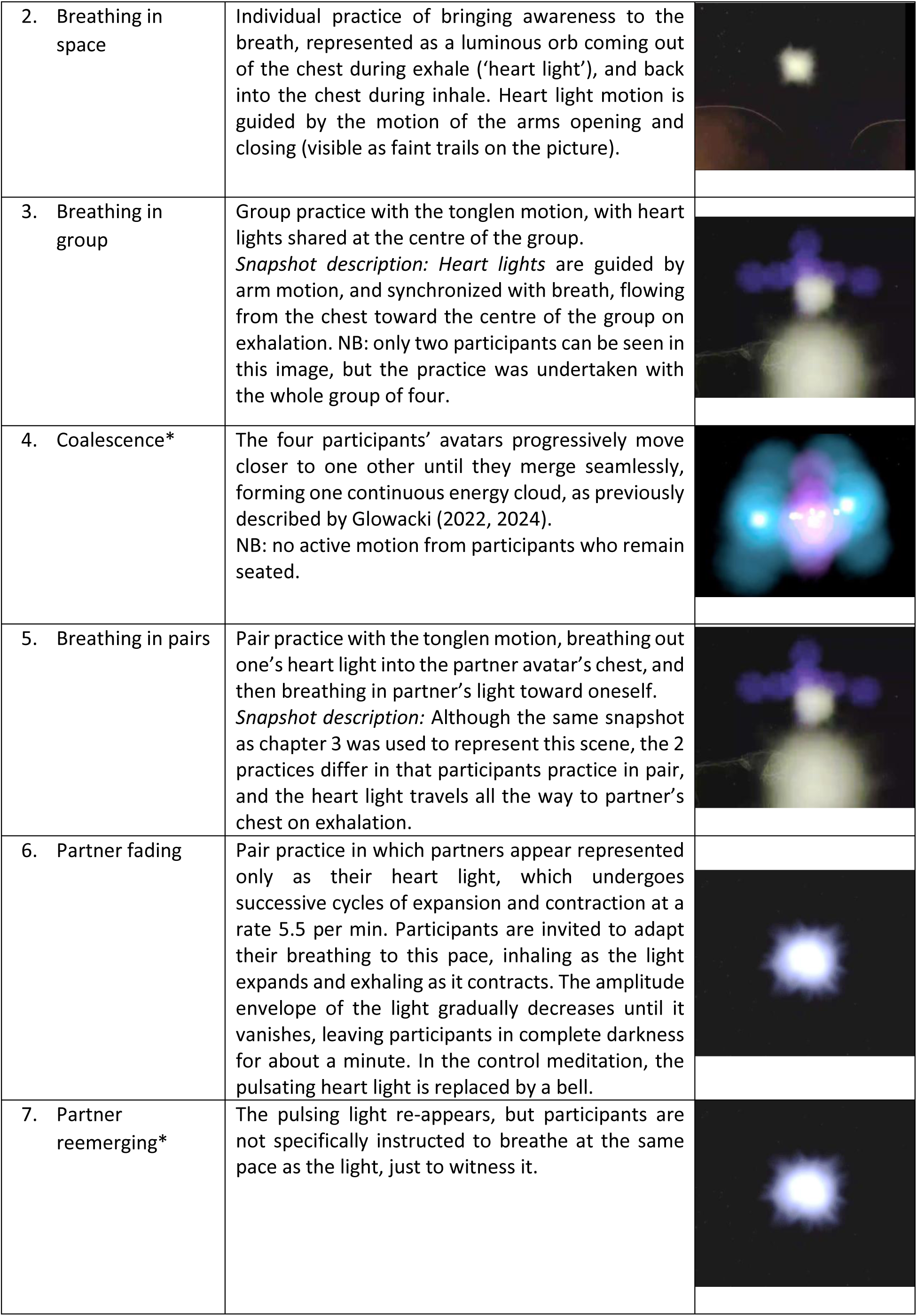
VR meditation chapters. **Table 1** provides a short description of each chapter in Ripple. The last column shows snapshots of each chapter, as seen from the point of view of a participant’s HMD. * indicates chapters that could be used for the physiological analysis (limited motion and natural breathing), see more details in method section ‘Chapters selected for physiological analysis’.

| Pre-recorded Ripple meditation chapters | VR meditation description | Snapshots |
| --- | --- | --- |
| 1. Mirror breathing | Pair practice involving arm motion (upward and downward) synchronized with slow box breathing, attuning to the breath and movement of partner.<br>*Snapshot description:* Partner's avatar with arms up. The 2 upper orbs represent hands, and the lower one represents the 'heart light'. |  |
| 2. Breathing in space | Individual practice of bringing awareness to the breath, represented as a luminous orb coming out of the chest during exhale ('heart light'), and back into the chest during inhale. Heart light motion is guided by the motion of the arms opening and closing (visible as faint trails on the picture). |  |
| 3. Breathing in group | Group practice with the tonglen motion, with heart lights shared at the centre of the group.<br>*Snapshot description:* Heart lights are guided by arm motion, and synchronized with breath, flowing from the chest toward the centre of the group on exhalation. NB: only two participants can be seen in this image, but the practice was undertaken with the whole group of four. |  |
| 4. Coalescence* | The four participants' avatars progressively move closer to one other until they merge seamlessly, forming one continuous energy cloud, as previously described by Glowacki (2022, 2024).<br>NB: no active motion from participants who remain seated. |  |
| 5. Breathing in pairs | Pair practice with the tonglen motion, breathing out one's heart light into the partner avatar's chest, and then breathing in partner's light toward oneself.<br>*Snapshot description:* Although the same snapshot as chapter 3 was used to represent this scene, the 2 practices differ in that participants practice in pair, and the heart light travels all the way to partner's chest on exhalation. |  |
| 6. Partner fading | Pair practice in which partners appear represented only as their heart light, which undergoes successive cycles of expansion and contraction at a rate 5.5 per min. Participants are invited to adapt their breathing to this pace, inhaling as the light expands and exhaling as it contracts. The amplitude envelope of the light gradually decreases until it vanishes, leaving participants in complete darkness for about a minute. In the control meditation, the pulsating heart light is replaced by a bell. |  |
| 7. Partner reemerging* | The pulsing light re-appears, but participants are not specifically instructed to breathe at the same pace as the light, just to witness it. |  |

### Description of the VR numadelic experience

The numadelic VR experience, called ‘Ripple’ was developed by two of the authors, David Glowacki and Justin Wall, who is a teacher in the Vajrayana Tibetan Buddhist tradition. This guided VR journey takes participants through successive practices (referred to below as ‘chapters’) involving breathing practices and playful exploration that invites participants to visualize their bodies as diffuse energetic essences (illustrated in Table 1). The audio file of the pre-recorded section of Ripple discussed in this article is available in the data repository, so are video snapshots for each chapter (see Data Availability).

Guided contemplative journeys within numadelic VR environments can support STEs by engaging several fundamental mechanisms underlying such states (Vidal et al., 2024). First, the aesthetics of the numadelic VR environment can enhance absorption, presence, and flow (Hooper et al., 2025; Magon and Cupchik, 2023; Suzuki et al., 2023). Second, embodying an ethereal ‘energy body’ within the VR space can foster greater permeability of self-boundaries, diminishing the rigidity of the embodied self and opening pathways to experiencing the properties of the VR avatar as one’s own - feeling more spacious, fluid and open, an effect which has been referred to as the ‘Proteus effect’ (Beaufils and Berland, 2022; Dupraz et al., 2024). Third, the multi-person aspect of the VR environment evokes a ceremonial space, priming participants for the emergence of uncommon emotional and cognitive states, and facilitating the dissolution of individual boundaries (Johnson et al., 1995; Rappaport, 1999; Whitehouse, 2002). Fourth, the group cohesion effect cultivated within this space can promote authentic dynamics of ‘interbeing’ and intimacy, nurturing a sense of group-directed transcendence. By creating a powerful sense of unity and shared experience, it dissolves the perception of self as an isolated entity, deepening the experience of interconnectedness (Dein, 2020). Lastly, bringing awareness to the breath is at the base of most meditation practices, and in numadelic VR, this process is facilitated by rendering the flow of breath visible through the mediation of the hand controllers. By gently moving their arms in what we refer to as the ‘tonglen motion’, expanding them outward (away from their chest) during exhalation and inward (toward their chest) during inhalation, in pace with their breathing, participants are enabled to visualize their breath in VR as a kind of proxy for live biofeedback using respiratory monitoring devices. This breathing motion also plays a crucial role in strengthening the connection between the user and their virtual avatar, enhancing the sense of immersion and presence. The breathing practices in pairs or in group also contribute to building group connection through physiological synchrony and a sense of cohesion between participants.

### Control meditation

The control meditation attempted to replicate each of these chapters outside VR, with participants being invited to close their eyes and visualise their body as the VR avatar – a cloud of energy, like a mist, without substance or solidity - and to imagine the other participants in a similar way. They were then guided through the same chapters as the VR session and invited to visualise with closed eyes what participants in the VR session see.

### Questionnaires

Trait questionnaires (baseline only):

- Sussex-Oxford Compassion for Others Scale (SOCS-O; Gu et al., 2020). It is a 20-item scale that measures the level of compassion toward others. It comprises five dimensions: 1) *Recognizing suffering* (e.g., "I notice when others are feeling distressed"), 2) *Understanding the universality of suffering* (e.g., "I understand that everyone experiences suffering at some point in their lives"), 3) *Empathizing with the person who is suffering* (e.g., "When someone is going through a difficult time, I feel kindly towards them"), 4) *Tolerating uncomfortable feelings* (e.g., "I stay with and listen to other people when they’re upset even if it’s hard to bear"), and 5) *Motivation to act* in order to relieve the other person’s suffering (e.g., "When others are struggling, I try to do things that would be helpful"). It uses a Likert-type scale ranging from 1 (never true) to 5 (always true).

State questionnaires (baseline and post session):

- The Hedonic and Arousal Affect Scale (HAAS; Roca et al., 2023) is a 24-item questionnaire designed to assess self-reported affective experiences, specifically focusing on the dimensions of valence (pleasantness/unpleasantness) and arousal (activation level). It employs a Likert-type scale ranging from 1 (not at all) to 4 (extremely).
- A Visual Analog Scale (VAS) to measure compassion and judgement towards others ad-hoc was developed from the Self-Compassion and Self-Criticism Scales (SCCS, Falconer et al., 2015). It uses a Likert-type scale ranging from 1 (not at all) to 7 (very much).
- A VAS was also used to measure connection to the session group (‘Connection to group’). It employs a Likert-type scale ranging from 1 (I feel completely disconnected) to 10 (I feel completely connected).
- Meditation pictographic scale (D’elia et al., 2025): a novel self-report tool designed to assess distinct meditative experiences through pictographic representations. The scale was initially developed with 20 items and later expanded to 23 items. Altogether, findings suggest the MPS offers a fair representation of meditation experiences, and that pictographic items help to provide more precise and unbiased measurement of these experiences. Items included in this study are those relevant to mood, relationality and self-transcendence: Positive affect, Negative affect, Compassion, Connection to others, perceived self-expansion and perceived porosity of body boundaries.

Additional questionnaire about the meditation experience (post session only):

- The Mystical Experience Questionnaire (MEQ-30; Maclean et al., 2012) is comprised of 30 items asking participants to reflect on their experiences of unity with the ultimate reality (Unity), feelings of ecstasy and peace (Positive Mood), perceptions of timelessness and loss of the sense of time (Transcendence of Time and Space), and the indescribable nature of their experience (Ineffability). It uses a Likert-type scale ranging from 1 (not at all) to 5 (extremely).
- The Awe Scale (Carrasco et al., 2018) is an instrument originally developed in Spanish for measuring state awe. It has 16 items classified in five factors: appraisals, affective response, physiological response, cognitive-subjective response and action tendency. It employs a Likert-type scale ranging from 1 (nothing) to 7 (a lot).

Subjective experience measures:

After their session, participants rated the quality of their experience along six different dimensions of interest: *Engagement*, *Valence*, *Intensity*, *Boundarilessness, Connection* and *Ego reduction* (Supplementary Table 1), focusing on each chapter separately. Before rating their experience on each chapter, they visualized a short video clip of the chapter to facilitate their recollection of the chapters (Table 1).

### Physiological data collection

#### Data collection

Cardiac and respiratory activity were recorded with the Shimmer 3 ECG module (Shimmer Sensing, Dublin, Ireland) with a sampling frequency of 180Hz. This wireless medical-grade chest electrocardiogram (ECG) monitor can simultaneously record ECG and impedance pneumography - which varies with each inhalation and exhalation - via four adhesive electrodes placed on the upper side of the chest and lower side of the abdomen. Electrodes were connected via four leads to the shimmer devices, which were strapped on a belt around participants’ abdomen. The four shimmer devices were wirelessly connected to the recording PC via Bluetooth, enabling data streaming and real-time monitoring, as well as pre-processing and extraction using the Consensys Pro software (v1.2.0, Shimmer).

#### Data preprocessing

ECG signal was first band pass filtered in Consensys (120 taps, cut off frequency of 0.5-30Hz) and saved for further pre-processing and analyses in Python using the Neurokit2 toolbox (Makowski et al., 2021). The function ecg_process was used for cleaning the data, detecting R peaks and calculating heart rate and RR intervals (time difference between two consecutive R peaks). To improve R peak detection, we applied the ‘kubios’ method (one of the methods options provided in Neurokit2) using the function ‘signal_fixpeaks’ which helps identify ectopic peaks. Data was visually inspected, and segments with poor R peak detection exceeding 10% of the analysis’s windows were excluded from analysis. One ECG dataset had to be excluded in the VR group, and three in the control group.

ECG-derived Measures of interest were computed as timeseries, and then averaged across the chapters of interest. These were:

##### Root Mean Square of Successive Differences (RMSSD)

RMSSD, a well-established measure of PNS activity that can be computed based on short data segments (minimun 10s) was calculated from RR intervals timeseries using 30s sliding windows, with 1s steps (Shaffer and Ginsberg, 2017).

##### Sympathetic and Parasympathetic indexes (SAI and PAI)

SAI and PAI calculation was performed following the methodology described in Valenza et al.(2018a), by uploading the timing of R events to the URL http://www.saipai-hrv.com/. These output series are derived from a Kalman-based estimation (Valenza et al., 2018b).

##### HRV amplitude

We propose a novel approach to capture dynamic sympathovagal engagement, derived from the amplitude of HR fluctuations over time, which we refer to as HRV amplitude (Figure 1). Unlike conventional measures or the related Local Power metric (which computes differences between local minima and maxima within each cardiac cycle and is highly sensitive to rapid, phasic PNS modulation) (Bornemann et al., 2016), our envelope-based approach provides a smoothed upper and lower bound of the HR signal over time. This smoothing attenuates ultra-fast parasympathetic dynamics, allowing the measure to reflect broader, sustained shifts in cardiac flexibility. Low HRV amplitude may occur either when SNS activity dominates PNS activity, or when both branches are minimally active, resulting in limited beat-to-beat variability. High HRV amplitude, on the other hand, reflects strong PNS modulation of cardiac activity associated with dynamic sympathetic counterbalance (i.e. the SNS is either co-active or in a state of high responsiveness). This dual dependence, highlighting HRV as a measure of autonomic modulation rather than PNS activity alone, reflects ANS’s branches dynamic regulation and interplay.

The amplitude of the RR interval signal envelope was computed at each time point by taking the difference between the moving median of local maxima and the moving median of local minima (Figure 1). This method, which uses a window of five data points, is robust to outliers, preserves local trends, and provides a stable estimate without excessive fluctuations. We used this approach rather than a Hilbert Transform because visual inspection revealed superior performance for highly entropic RR interval time series.

Since conditions varied throughout the meditation session, and given our focus on a measure that may ultimately reflect real-time subjective experience, we did not compute HRV parameters requiring longer data windows for reliable estimation - such as frequency-based and non-linear metrics.

##### Respiratory Rate & Amplitude

The guided meditation, both in VR and audio format, involved various slow breathing exercises (see Table 1) which are likely to affect HRV measure. Intentional regulation of breathing is one of the most direct ways to influence autonomic activity, since inhalation is linked to SNS and exhalation to PNS activity (Fincham et al., 2023). This relationship is referred to as respiratory sinus arrhythmia, where HR rises during inhalation and falls during exhalation, and is reflected in the HRV around frequencies corresponding to breathing rate (∼ 0.12 to 0.4Hz) (Eckberg, 1983; Menuet et al., 2025). In addition, excessive lung inflation during forced slow breathing activates the Hering–Breuer reflex, which halts inspiration and can affect HRV (Quigley et al., 2024). Respiratory signal was therefore monitored to account for breathing patterns and the way they influenced HRV, and chapters where participants were specifically instructed to breathe other than naturally were not included in the HRV analysis.

Respiratory signal was low pass filtered in Consensys (120 taps, cut-off frequency of 3Hz) before being exported for further pre-processing in Python using the rsp_process function from the Neurokit2 toolbox (Makowski et al., 2021). The raw signal, sampled at 180 Hz, was then further preprocessed using a second order 0.05-3 Hz bandpass Butterworth filter (Khodadad et al., 2018). The peak detection was carried out using the method described in Khoadadad et al. (2018). To ensure data quality, visual inspection was performed using the same approach as for ECG signals. Datasets were excluded if artifacts or detection issues exceeded acceptable thresholds, leading to the removal of 2 respiratory datasets in the VR group and 5 in the Control group. Finally, respiratory rate and amplitude were extracted from the cleaned respiratory signal for further analysis.

### Chapters selected for physiological analysis

The VR experience *Ripple* was designed independently of this study and it did not incorporate design considerations for consistent HRV monitoring. While we introduced modifications (e.g., participants were seated rather than standing), the experience inherently includes slow breathing practices and arm motions that substantially influence HRV measurements (Quigley et al., 2024). We reasoned that these physiological manipulations could introduce noise into the relationship between HRV indices and subjective reports. To minimize these confounds, our analysis focused exclusively on Chapters 4 and 7, which are the only segments without guided respiration or arm movements. These chapters notably correspond to the highest reported levels of self-transcendence (see Figure 3, *Ego Reduction*). To enhance statistical power and reduce the number of statistical tests, data from these two chapters were averaged prior to correlation analyses, though separate results are also reported.

**Figure 3:**
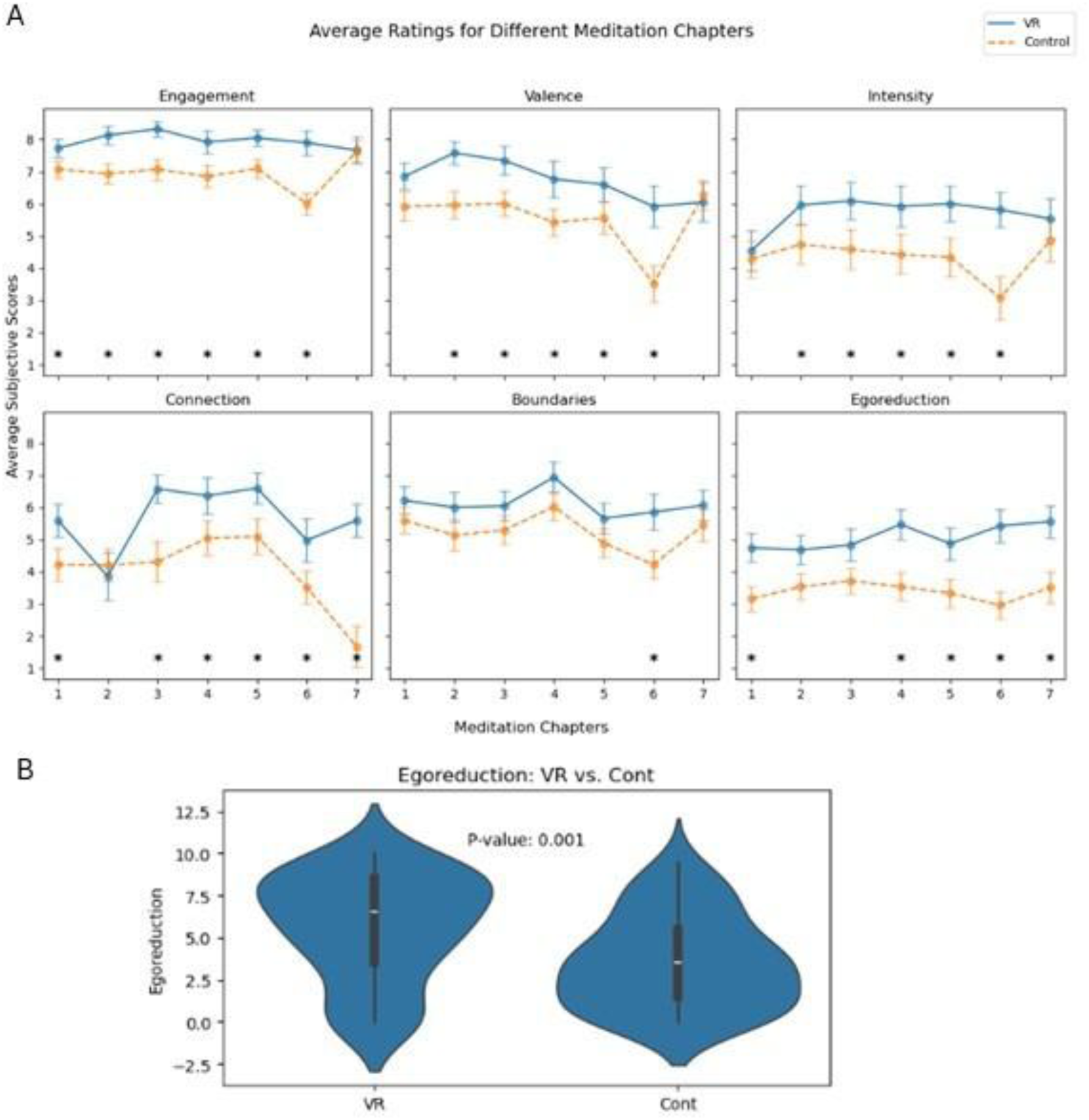
A) Averaged subjective ratings for each of the 7 chapters of the VR (continuous blue line) and control (dotted orange line) guided meditations. * indicates significant differences between groups (p<0.05, Mann-Whitney U test). B) Violin plots illustrating the difference in Ego reduction ratings between participants in the VR and Control groups, averaged across Chapters 4 and 7. The p value relates to the statistical comparison between the two groups using a Mann-Whitney U test.

### Statistical analysis

Data normality was assessed using the Shapiro-Wilk test. When comparing VR and Control groups, normally distributed variables were analyzed using an independent t-test, while non-normally distributed variables were assessed using the Mann-Whitney U test.

To examine the relationship between physiological and psychological variables within groups, all variables were normalized using a min-max scaler, and Spearman correlations were performed, controlling for Age, Gender, and BMI as covariates.

Given our hypothesis-driven approach with a primary focus on HRV amplitude in relation to STEs, we did not perform multiple comparison corrections on the corresponding correlation. Other correlations reported for comparative purposes were likewise not corrected, serving as reference points for HRV amplitude performance relative to other measures.

A piecewise linear regression (also known as segmented regression) approach was used to model the relationship between *Ego reduction* (independent variable) and HRV amplitude (dependent variable), allowing for the identification of a naturally occurring threshold or breakpoint in the data. The optimal breakpoint location was first estimated using non-linear optimization techniques implemented in the ‘piecewise-regression’ Python library. To control for potential confounding factors, a multiple linear regression model was subsequently implemented using the ‘statsmodels’ library. This model included the primary independent variable, a generated indicator variable for the determined breakpoint, and covariates including BMI, age, and gender. The statistical significance of the breakpoint was assessed using a pseudo score test (Davies test), and the overall significance of predictors was determined using standard p-values from the OLS regression summary, with a significance threshold set at *α*=0.05.

To explore whether HRV amplitude’s relationship with psychological measures was mediated by other physiological metrics (notably respiratory amplitude), we employed partial correlations and mediation analysis, using Pingouin’s toolbox in Python (Vallat, 2018).

Participants’ psycho-emotional state was assessed before and after the session through multiple measures, grouped into three domains to streamline analysis: 1) Positive Affect – Pictographic scale (Positive affect), HAAS (Positive valence emotions), 2) Negative Affect – Pictographic scale (Negative affect), HAAS (Negative valence emotions), and 3) Relationality – Pictographic scales (Compassion, Connection group, Connection others). False Discovery Rate (FDR) correction was employed to correct for multiple comparison.

### Analysis of DMT Data

In order to evaluate the robustness and generalizability of our results, and test whether the autonomic signature observed in numadelic VR (specifically HRV amplitude) also emerges in a pharmacologically induced context, we revisited data from a previously published study examining the impact of DMT on ANS activity (Bonnelle et al., 2024). The same ECG data processing protocol as described above was employed. Out of the 17 datasets used in the original study, 4 had to be excluded due to noisy signals that could not be adequately corrected using our automated processing pipeline.

For each participant, ECG data consisted of 8 minutes of baseline recording, followed by 20 minutes of post-injection data. HRV amplitude was computed as a time series across the entire session and then averaged after the initial physiological stress response induced by DMT injection, starting at minute 4 post-injection and continuing until the end of data collection.

These averaged HRV amplitude measures were then correlated using Spearman correlations with subscales from two questionnaires assessing the quality of peak experiences: the Mystical Experience Questionnaire (MEQ-30) and the 11 Dimensions Altered States of Consciousness Questionnaire (11D-ASC). The 11D-ASC features the following subscales: Experience of Unity, Spiritual Experience, Blissful State, and Insightfulness Impaired Control and Cognition, Anxiety Disembodiment, Complex Imagery, Elementary imagery, Synesthesia and Meaning (Studerus et al., 2010). False Discovery Rate (FDR) correction was employed to correct for multiple comparison.

## Results

### Subjective experience during Numadelic VR meditation compared to control meditation

Figure 3 illustrates the differences observed between the VR and Control groups across the six different subjective experience measures, and throughout each of the seven chapters. *Engagement*, *Valence* and *Intensity* of emotions were significantly higher during VR on most chapters, apart from the first and last one (p<0.05). *Connection* and *Ego reduction*, two measures reflecting self-transcendence (Yaden et al., 2017), were also significantly higher in the VR group on chapters 1, 4, 5 6 and 7, and also chapter 3 for connection (p<0.05), consistent with previous findings that numadelic experiences can achieve a significant degree of self-transcendence (Glowacki et al., 2022; Vidal et al., 2024). When averaged across chapters 4 and 7, on which we focus our analysis, the VR group (mean =5.64) shows significantly higher ratings *than the control group (mean=3.54) on Ego Reduction,* the subjective rating best reflecting STE (U = 1405.5, z = 3.17, p = 0.0015). Surprisingly, however, the perceived *Boundaries* scores did not statistically significantly differ between groups, possibly the result of participants in the Control group having their eyes closed and being explicitly invited to visualise their body as a cloud without substance or solidity.

### Relation between STE ratings and HRV amplitude during the numadelic experience

Both motion and breathing can significantly influence ANS activity, and failing to account for them can confound the relationship between HRV and subjective measures of emotion (Quigley et al., 2024; Shaffer and Ginsberg, 2017). To minimize these confounding effects, we focused our analysis on Chapters 4 (Coalescence) and 7 (Re-emergence), during which participants experienced neither imposed breathing nor arm motion. Incidentally, these two chapters also correspond to the peak phases of the experience, characterized by high Ego reduction ratings (Fig. 3).

As hypothesized, HRV amplitude on both chapters showed a positive correlation with *Ego reduction* ratings (Partial Spearman correlation controlling for Age, gender and BMI, Chapter 4: n = 44, ρ = 0.36; CI95% = [0.05, 0.60], p = 0.02, Chapter 7: n = 44, ρ = 0.33, CI95% = [0.02, 0.58], p = 0.03). The consistency of this relationship suggests that both segments capture the same underlying physiological correlate of the subjective experience. Despite the lack of homogeneity in HRV amplitude distribution in chapters 4 and 7 (see Table S2), and given the rationale behind selecting these two chapters, free of externally imposed confounds, we proceeded with the aggregation of data from these two chapters for the subsequent analyses, rather than reporting results separately for each one, thereby enhancing statistical power and providing a more stable estimate of autonomic function during peak STE. The correlation between *Ego reduction* and *HRV amplitude* remained after aggregating the two chapters (n = 44, ρ = 0.47, CI95% = [0.19, 0.68], p = 0.001, Figure 4A).

**Figure 4:**
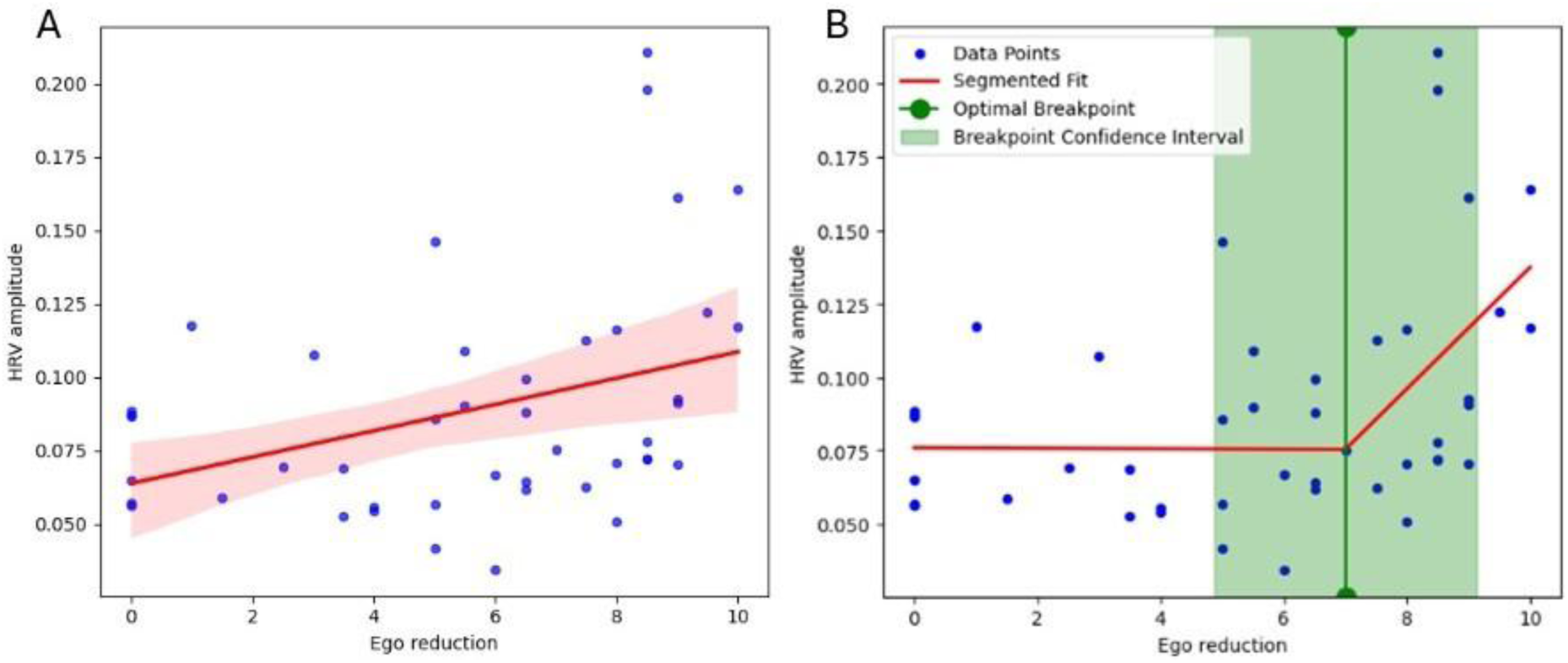
Comparison of a linear regression (A) and segmented regression (B) modeling HRV amplitude in relation to Ego reduction Ratings. Regression lines are shown in red, with the surrounding pink area indicating the 95% confidence interval on the left panel. The green vertical line on the right marks the optimal breakpoint identified in the segmented fit, and the shaded green band denotes its 95% confidence interval.

Visual inspection suggests a potential inflection in the relationship between Ego reduction and HRV amplitude. To examine this in more detail, a piecewise linear regression model was fitted, while statistically controlling for BMI, age, and gender using multiple ordinary least squares regression. The overall model was statistically significant (R2=0.270, F(5,41)=3.034, p=0.020). The Davies test confirmed that the segmented model provided a significantly better fit than a single linear regression (p=0.029). The analysis revealed a significant breakpoint at an ego reduction score of 7.0 (confidence interval [4.9, 9.1]). Below this threshold, there was no significant relationship observed between Ego reduction and HRV amplitude (*β*=−0.0002, p=0.952). However, for Ego reduction scores above the breakpoint, a significant positive association emerged (*βc*ombined=0.0218, 95% CI [0.002, 0.041], p=0.029), indicating that higher levels of ego reduction beyond the threshold correlated with increased HRV amplitude. Of the covariates included, gender (p=0.948) and BMI (p=0.553) did not show a significant association with HRV amplitude, while age was marginally non-significant (p=0.081).

To evaluate how HRV amplitude compares to other autonomic activity measures in capturing STEs during the VR session, we report Spearman correlation coefficients between autonomic metrics (HR, RMSSD, PAI, SAI, respiratory amplitude and respiratory rate) and subjective ratings (Figure 5). Since respiratory parameters were included in this analysis, only participants whose respiratory data could be reliably analyzed were considered (n = 40). Within this subset, HRV amplitude remained significantly correlated with *Ego reduction* ratings (ρ = 0.47, CI95% = [0.18, 0.69], p = 0.003). It was also marginally correlated with *Connection* ratings (ρ = 0.30, CI95% = [-0.04, 0.55], p = 0.08). Respiratory amplitude was the only other measure showing a correlation with *Ego reduction* ratings (ρ = 0.40, CI95% = [0.09, 0.64], p = 0.01).

**Figure 5:**
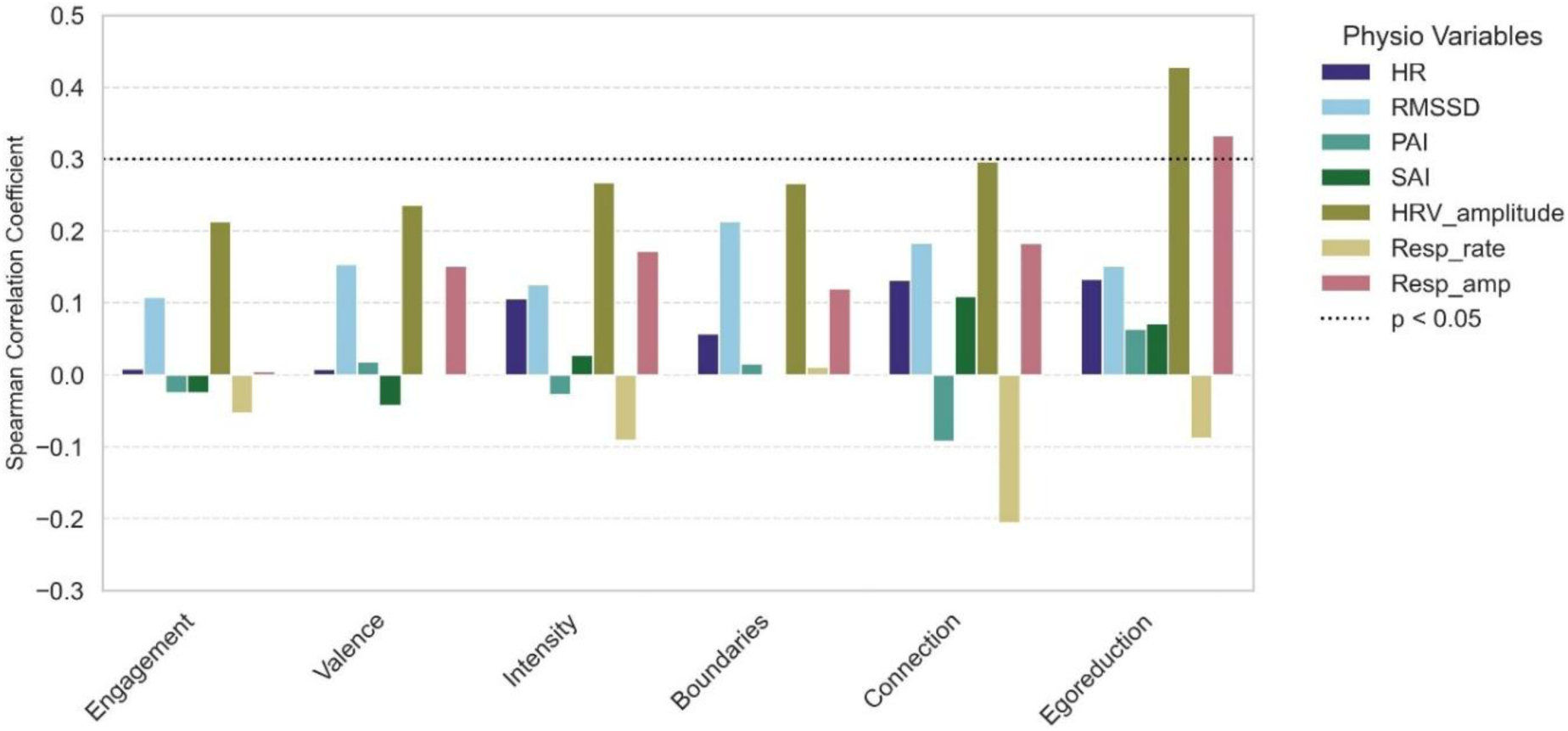
Spearman linear correlation coefficients for each correlation between the six subjective ratings (horizontal axis) and the 8 physiological variables in the VR group, for participants with reliable ECG and respiratory data (n=40). The dotted line indicates the significance threshold, so that coefficients above this threshold correspond to statistically significant correlations.

To explore the possibility that respiratory amplitude may mediate the relation between HRV amplitude and Ego reduction ratings, we performed a mediation analysis controlling for age, gender, and BMI, with normalized data (z-scoring). Given the non-normal distribution of the data, a bootstrapped approach with 5,000 resamples was employed to estimate confidence intervals. Results indicate that respiratory amplitude did not significantly mediate the relationship between *HRV amplitude* and *Ego reduction* (Indirect effect size = 0.06, Bootstrapped 95% CI: [-0.13, 0.25]). The direct effect of HRV amplitude on *Ego reduction*, adjusted for covariates, was 0.35 (95% CI [–0.01, 0.72]), suggesting a positive but statistically inconclusive relationship. This relatively wide confidence interval reflects an apparent instability in the direct path that could be explained by the complex relationship between HRV amplitude and *Ego reduction*, which may have been underestimated or obscured by the linear mediation model.

Finally, to assess whether baseline *HRV amplitude* might predispose individuals to stronger STEs, we correlated baseline *HRV amplitude* with average *Ego reduction* ratings during the session. This association was not significant (ρ = –0.07, p = 0.84), suggesting that HRV amplitude does not represent a stable trait or inherent propensity to experience STEs. Instead, it appears to reflect a dynamic state of autonomic function expressed during the experience itself.

### Relation between HRV amplitude and STE ratings during the control meditation

Based on the observed nonlinear relationship between HRV amplitude and *Ego reduction* in the VR group, and given that *Ego reduction* ratings were significantly lower in the Control group, we anticipated that the association between HRV amplitude and *Ego reduction* in this group would be weaker or potentially absent. This expectation was confirmed by the lack of a significant correlation between *Ego reduction* ratings and HRV amplitude in the Control group (n=42, ρ=-0.07, p= 0.65). Within this group, no significant relationship was detected between physiological and subjective measures.

### Relation between HRV amplitude during the VR experience and changes in psycho-emotional state after the experience

Averaged HRV amplitude during Chapters 4 and 7 of the VR session significantly and positively correlated with changes in *Positive affect* (n=44, ρ=0.43, CI95%=[0.14, 0.65], p= 0.015) and negatively with changes in *Negative affect* (n=44, ρ=-0.39, CI95%=[-0.63, −0.1], p=0.015) after the session relative to baseline, but not with changes in *Relationality* (ρ=0.26, p=0.10). P values are FDR corrected for multiple comparison.

When looking at the impact of other autonomic measures, respiration amplitude during the session also appeared to positively impact *Positive affect* (ρ=0.41, CI95%=[0.11, 0.64], p= 0.03), as well as *Relationality* (ρ=0.35, CI95%=[0.04, 0.6], p=0.045), but not *Negative affect* (ρ=-0.21, p=0.18). P values are FDR corrected for multiple comparison. Average measures of Positive affect, Negative affect and Relationality for baseline and post-session are reported on Table S3.

### Relation between HRV amplitude and STE questionnaires

Two questionnaires assessing participant’s STE were used: the MEQ-30 and the Awe Scale. They were administered after the session, with respect to the entire session (rather than specific chapters). None of these scales showed significant correlation with *HRV amplitude* (MEQ-30_Tot: ρ=0.13, p=0.41; Awe_Tot: ρ=0.18, p=0.24), although they were both correlated with subjective ratings, particularly robustly with *Ego reduction* (MEQ: ρ=0.73, Awe: ρ=0.75, p<0.001) and *Boundaries reduction* (MEQ: ρ=0.77, Awe: ρ =0.73, p<0.001). None of the other physiological measures correlated with these questionnaires.

### Relation between Compassion traits, quality of the experience and HRV amplitude

Higher levels of compassion trait, as measured by the SOCS-other questionnaire at baseline, were associated with higher ratings on all the subjective ratings dimensions during the VR meditation, particularly *Ego-reduction* (ρ=0.47, p<0.001), and *Connection* (ρ=0.43, p=0.002). Higher compassion trait was also associated with a more pronounced increase in *Relationality* after the experience compared to baseline (ρ=0.33, p=0.025).

Importantly, compassion traits were also associated with significantly higher *HRV amplitude* during the VR experience (SOCS for others (total score) n=44, ρ=0.50, CI95%=[0.23, 0.70], p<0.001). The SOCS subscales with the strongest influence on this relationship were ‘*Recognizing suffering*’ (ρ=0.51), ‘Feeling for others’ (ρ=0.41), and ‘*Tolerate suffering*’ (ρ=0.39). The subscales ‘*Understanding the universality of suffering*’ (ρ=0.06) and ‘*Motivation to act*’ (ρ=0.27) did not significantly correlate with HRV amplitude. However, there was no significant relationship between Compassion traits and HRV amplitude at baseline.

*RMSSD* (n=44, ρ=0.44, p=0.004) and *Respiratory amplitude* (n=0.40, ρ=0.43, p=0.006) during the VR experience were also positively correlated with compassion traits at baseline.

### Assessing HRV Amplitude as a Physiological Marker of Self-Transcendence in Psychedelic States

To evaluate the effectiveness of HRV amplitude in capturing STEs within other NOSC, we revisited previously published ECG data collected during DMT-induced STEs. First, we averaged across each participant HRV amplitude time course in order to visualise how HRV amplitude fluctuates, on average, throughout the entire DMT experience. As illustrated on Figure 6, HRV amplitude increases approximately 10 minutes post-injection, possibly reflecting the previously described window of sympathovagal engagement associated with vagal rebound following the initial peak of sympathetic activity induced by DMT, which we refer to as the ‘post-acute stress response’ phase (Bonnelle et al., 2024).

**Figure 6:**
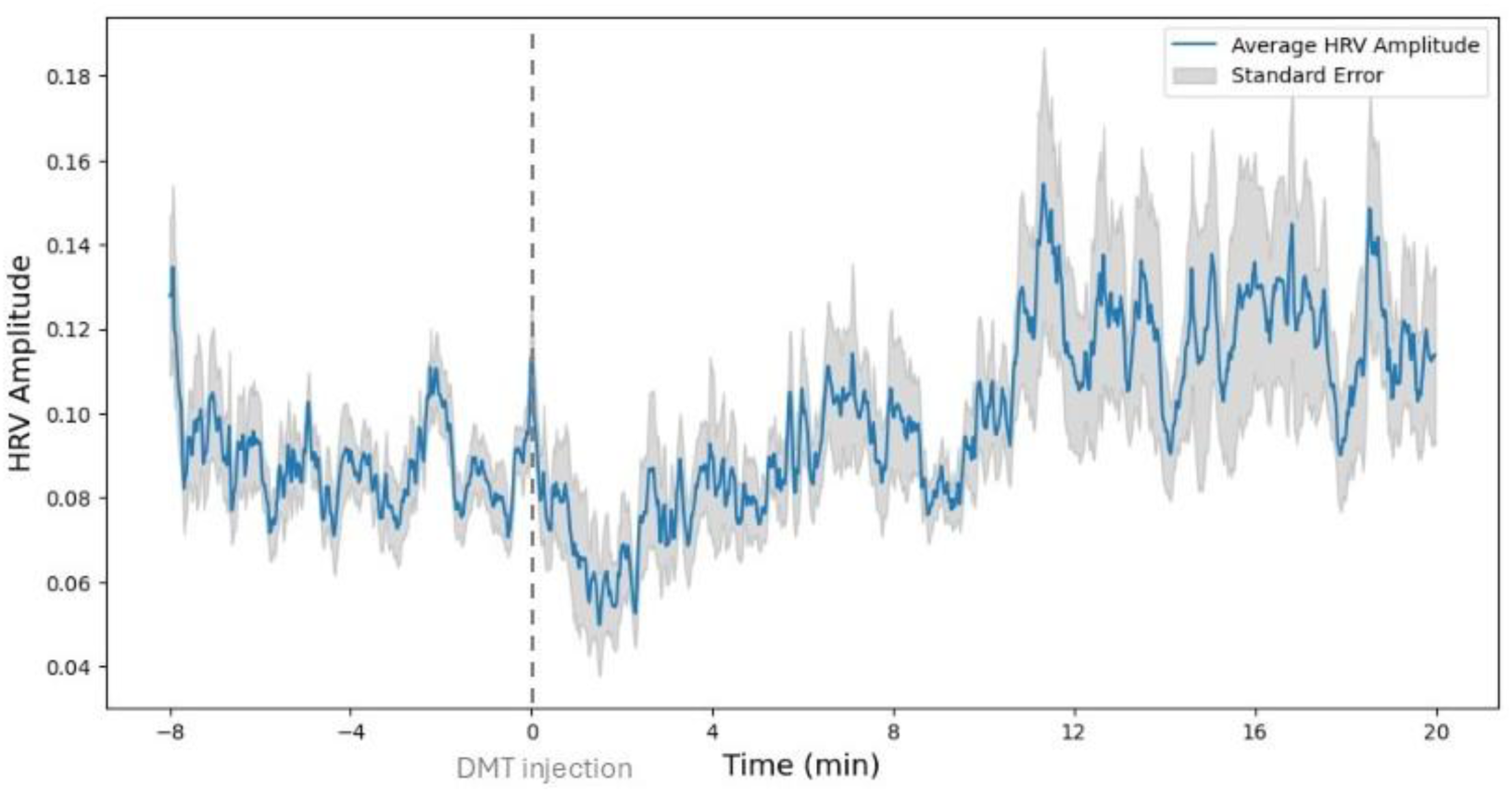
Fluctuations in HRV amplitude averaged across 13 participants. The grey area indicates standard errors. DMT was injected following a 8-min baseline period (grey dotted line).

For each participant, HRV amplitude was averaged across a 4–20 min post-injection window, corresponding to the post-acute stress response phase. This choice was guided by our vagal rebound hypothesis and consistent with our prior DMT study, which identified an initial sympathetic surge that typically resolves within ∼3 min post-injection (Bonnelle et al., 2024). Averaging from minute 4 onward therefore captures the more stable physiological period of the session.

HRV amplitude measures averaged over this period were correlated with subjective ratings of peak experience collected with the Mystical Experience Questionnaire (MEQ30) and the Altered State of Consciousness scale (11D-ASC). Average scores on the MEQ-30 where robustly correlated with *HRV amplitude* (N=13, spearman ρ =0.82, CI95%=[0.50, 0.93], p=0.001), which was mostly driven by the two subscales ‘*Mystical experience*’ (ρ =0.77, CI95%=[0.43, 0.94], p=0.003), and ‘*Positive mood*’ (ρ =0.73, CI95%=[0.34, 0.93] p=0.005). In addition, HRV amplitude was also highly correlated with the 11D-ASC dimension ‘*Blissful state*’ (ρ=0.80, CI95%=[0.43, 0.94], p=0.001), and more marginally, with the subscale *‘Oceanic Boundlessness*’ (ρ =0.55, CI95%=[-0.045, 0.84], p=0.05). P values are FDR corrected for multiple comparison. Correlations can be visualised on Figure 7.

**Figure 7:**
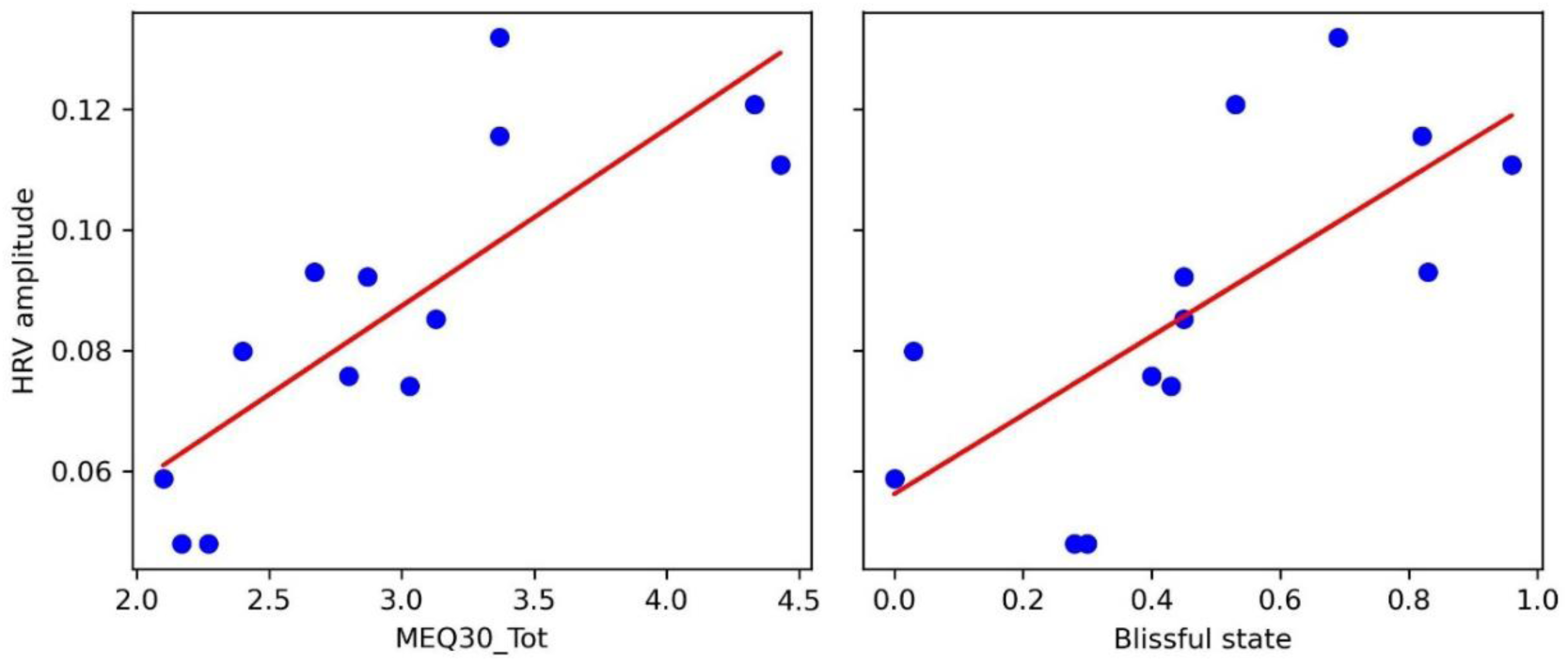
Correlations between HRV amplitude during the post-acute stress response phase following DMT injection, and ratings of STEs collected with the MEQ-30 and the 11d-ASC (Blissful state).

Although the sample size for the DMT dataset was relatively small (N = 13), which warrants caution in interpretation, the correlations between HRV amplitude and MEQ-30/11D-ASC scores were based on a priori hypotheses and, in both cases, appear visually well supported by the spread of data points along the linear regression lines (Figure 7). Moreover, the use of Spearman correlations mitigates the potential impact of outliers. These considerations suggest that the observed associations are unlikely to be driven by individual cases, though replication in larger samples will be needed.

## Discussion

The central question driving this study was if STEs experienced during numadelic VR may be captured at the physiological level using a measure reflecting sympathovagal engagement. Addressing this is essential for advancing research into these NOSC, without solely relying on participants’ subjective reports, which are limited in time resolution, labor-intensive to collect, and dependent on accurate recollection and interpretation of experiences along specific questionnaire dimensions. Building on prior research into the role of the ANS in psychedelic peak experiences, we introduce a novel measure, referred to as HRV amplitude, to capture sympathovagal influence on cardiac activity. Consistent with our hypothesis, findings indicate that HRV amplitude significantly correlates with STE ratings across both numadelic and psychedelic-induced STEs. In addition, this measure is also related to compassion traits, and reflects the positive after-effects of numadelic sessions on participants’ emotional state.

### Numadelic VR as a promising tool to study STEs

Research on STEs has been constrained by their inherently non-ordinary nature, as these states typically require years of advanced contemplative practice or the use of mind-altering substances like psychedelics. The potential of numadelic experiences to induce STEs in individuals is remarkable and holds significant value not only for research, but also for therapeutic applications and personal development. Amid the psychedelic research renaissance, one of the most consistent themes emerging over the past decade is the central role of STEs in the long-term benefits of psychedelic-assisted therapy (Ko et al., 2022). As modern psychiatry increasingly acknowledges the therapeutic potential of these states for mental health, there is growing interest in developing novel, safe, and reliable methods to induce such experiences, alongside tools to monitor their occurrence and intensity.

Our results indicate that numadelic VR is not only more engaging than an audio-guided meditation, but also more effective at lowering the sense of ego, while fostering connection between participants. As outlined in the Methods section, several core features of numadelic experiences contribute to their potential for STEs: 1) Greater absorption, fostering deep immersion and altered attentional states; 2) Aesthetic qualities characterized by low representational imagery, evoking the essential traits of STEs (non-rigidity, openness, and shifts in self-perspective); 3) Ceremonial space, reinforcing meaning and engagement and 4) Social connection, enhancing intersubjective experiences. Together, these features reduce cognitive rigidity, enhance openness, and encourage altered self-perception, making them potent facilitators of STEs.

Another potentially relevant component underlying the impact of numadelic VR may be summarized as ‘safe exposure’. VR has the potential to create a structured yet fluid environment where individuals may feel safe enough to lower their defences and explore altered states of consciousness. Much like exposure therapy, where a controlled space allows patients to gradually confront their fears without overwhelming anxiety (Boeldt et al., 2019), a numadelic-guided experience may act as a container that enables individuals to let go of rigid identity structures, allowing for ego dissolution, self-transcendence, or profound shifts in awareness.

### Physiological marker of STEs

Previous research on the physiological basis of STEs has largely focused on their neurophysiological correlates. While neural activity provides valuable insights into the phenomenology of these experiences, its measurement is often costly, computationally demanding, and impractical outside laboratory settings. In contrast, moment-to-moment cardiac metrics offer a more ecologically valid and accessible approach. Drawing from prior research on psychedelics (Bonnelle et al., 2024), where shifts in ANS function have been implicated in NOSC, we hypothesized that ANS activity could serve as a physiological marker of STEs. More specifically, we hypothesized that STEs manifest as a paradoxical state of ‘relaxed alertness’, in which both sympathetic and parasympathetic branches of the ANS may be co-activated. Since HRV amplitude reflects the dynamic interplay between these systems, it presents a promising moment-to-moment indicator of sympathovagal engagement. To assess its effectiveness, we compared this measure with other short-segment estimations of autonomic function. Although HRV amplitude partially correlated with other parasympathetic markers, it was the only cardiac-derived measure to significantly correlate with ego reduction ratings, suggesting that this relation was not mostly driven by the parasympathetic component of HRV amplitude.

Our results indicate that ego reduction is associated with higher HRV amplitude, but the relation does not appear to be bi-directional: higher HRV amplitude at baseline did not necessarily give rise to higher ego reduction ratings during the experience. HRV amplitude may therefore serve as a useful biomarker of NOSC, but may not be in itself a sufficient causal mechanism. Other elements must converge to produce such experiences, with autonomic state being only one component. At the CNS level, STEs are consistently linked to reductions in default mode network (DMN) activity across pharmacological and non-pharmacological contexts, supporting a temporary attenuation of self-related processing (Millière et al., 2018). The DMN’s activity is inversely correlated with the Central Executive Network (CEN) and the Salience Network (SN) (Menon and Uddin, 2010; Sridharan et al., 2008). During STE, the SN may act as a switch, down-regulating the DMN and ramping up the CEN and other task-relevant networks, leading to higher level of absorption and low self-referential thinking, also described in flow states (van der Linden et al., 2021).

The relation between HRV amplitude and ego reduction ratings was non-linear, so that the two variables were more related with each other when participants’ STE ratings were above a certain threshold. Possible interpretations of this observation are discussed below.

### ANS and emotions

Research on the relationship between emotional states and ANS activity suggests that the connection is complex, influenced by multiple modulating factors (Friedman, 2010; Kop et al., 2011; Kreibig, 2010). We propose that specific conditions must be met for individuals’ subjective experiences to reliably map onto their ANS activity.

Emotional influences on the ANS arise through both bottom-up and top-down processes. On the one hand, visceral and sensory inputs can trigger autonomic adjustments that accompany affective states; on the other, higher-order cortical and limbic regions exert central modulation, shaping autonomic output in line with appraisal, attention, and regulation (Schneider et al., 2025; Thayer and Lane, 2009, 2000). At the same time, it is important to recognize that a substantial portion of autonomic activity reflects homeostatic and survival functions (e.g. cardiovascular regulation, thermoregulation, digestion) that are not directly tied to emotion (Purves et al., 2001). Distinguishing emotion-related variance from these broader physiological functions is therefore essential for relating ANS measures to subjective experience (Kok and Fredrickson, 2010; Schneider et al., 2025). This relationship may strengthen with more intense emotional experiences but can become less distinguishable within autonomic signals for subtle or secondary emotions (Levenson, 2014). This could explain at least in part the presence of a ‘breakpoint’ in the relationship between HRV amplitude and ego reduction rating, after which the relation between the two variables becomes significant. Additionally, body motion and imposed breathing introduce variability that can diminish the portion of ANS variance relevant to emotion (Bernardi et al., 2001; Laborde et al., 2017; Norman et al., 2014). To minimize these confounding influences, our analysis focused on periods when participants remained relatively still and breathed naturally.

Secondly, the accuracy of subjective experience reports likely correlates with presence and absorption levels. At the neural level, high-absorption states, where individuals are deeply immersed without deliberate effort, are linked to bottom-up (stimulus-driven) attention, characterized by enhanced sensory and interoceptive processing (Barrett et al., 2004). This contrasts with, on the one hand, DMN activity, which prioritizes energy-saving low sensory engagement (Raichle, 2015), and on the other hand, top-down attentional control, which relies on prior expectations rather than real-time sensory integration (Corbetta and Shulman, 2002). When individuals experience heightened presence, they are therefore more likely to accurately report their internal state than those engaged in abstract cognition or mind-wandering.

Taken together, optimal conditions for aligning subjective experience with ANS activity may involve physical stillness, natural breathing, and heightened present-moment absorption - the foundation of most meditation practices, which aim to harmonize physiological and psychological states (Lutz et al., 2008; Tang et al., 2015). Consistent with this, our results indicate that STE ratings correlated with ANS activity during numadelic and psychedelic experiences, but not during a control meditation performed by non-advanced meditators, who reported significantly lower levels of ego reduction. This difference may reflect a greater body-mind disconnect in ordinary states of consciousness. Additionally, the non-linear relationship between STE ratings and HRV amplitude suggests that physiological markers become more reflective of self-transcendent states as experience depth increases.

### Respiratory measures

Our results indicate that respiratory amplitude plays a favorable role in the experience, showing a moderate correlation with ego reduction ratings during the session, though less robustly than HRV amplitude. Additionally, respiratory amplitude was associated with beneficial post-session effects on affect and relationality.

Despite both respiratory rate and respiratory amplitude being correlated with HRV amplitude (one negatively, the other positively), only respiratory amplitude was significantly linked to STE ratings during the VR session. A mediation analysis further revealed that the relationship between HRV amplitude and STE ratings was not mediated by respiratory amplitude, suggesting that HRV and respiratory amplitude contribute to STE through distinct pathways. While our mediation analysis did not support respiration as a mediator of the HRV/ego reduction relationship, respiration may nonetheless modulate the strength or shape of this association. Given the close physiological coupling between respiratory variability and HRV amplitude, particularly with slow breathing (Eckberg, 1983; Grossman and Taylor, 2007; Menuet et al., 2025), fluctuations in breathing could amplify or attenuate HRV responses, thereby influencing the observed correlation with ego reduction. This perspective is consistent with our finding of non-linear relationship, in which the positive association between HRV amplitude and ego reduction only becomes apparent above a threshold value. Respiratory variability may contribute to this non-linear pattern by interacting with autonomic dynamics, suggesting that modulation rather than mediation may provide a more physiologically coherent framing of the interaction between the two systems.

Slow and high amplitude breathing practices that engage the abdomen (diaphragmatic breathing) or the entire torso (Dirga pranayama, also known as Three-Part Breath or Complete Breath) are well-documented for their positive effect on emotional regulation (Hamasaki, 2020). At the physiological level, high-amplitude breathing increases carbon dioxide partial pressure in the blood, which enhances cerebral blood flow, optimizes oxygen delivery to the brain, and directly benefits cognitive function and emotional stability (Goheen et al., 2023). In addition, breath amplitude has been shown to entrain brainwave activity, potentially enhancing focus, emotional resilience, and stress regulation (Allen et al., 2023; Brændholt et al., 2023; Heck et al., 2017). Respiratory amplitude may thus serve as another, albeit weaker, physiological marker of STE in numadelic experiences. However, it is important to note that with currently available biosensors, recording high-quality respiratory signals is more challenging than HRV.

### HRV amplitude and compassion

Our results indicate that individuals with higher compassion traits exhibit higher HRV amplitude levels during the numadelic experience, consistent with prior work on the physiological correlates of compassion (Bornemann et al., 2016; Di Bello et al., 2020; Förster and Kanske, 2022; Stellar et al., 2017). This effect was not observed at baseline, suggesting that HRV amplitude reflects state-dependent (acute) compassion rather than a stable trait, which may be more closely tied to higher-order cognitive processes. Although participants may not have consciously experienced compassion during the VR session, the modulation of HRV appears to index a shared physiological mechanism linking compassion and self-transcendence, potentially through absorption, decentering and/or broadening of attentional scope (Förster and Kanske, 2022). Taken together, these findings suggest that HRV amplitude reflects flexible autonomic regulation underlying compassion, and in the numadelic context, this same flexibility may support self-transcendent states characterized by emotional openness, regulation, and engagement.

### Are all STEs associated with the same physiological state?

While STEs have only recently gained attention in modern science, they have long been explored in Eastern traditions, particularly through Buddhist jhanas—eight progressively deeper states of mental unification and absorption. Early jhanas are marked by sensory-rich experiences such as joy and energy, but as practice deepens, sensory content diminishes, giving way to equanimity and inner stability (Gunaratana, 1988).

The ANS involvement in these stages remains largely unexplored, though deeper states are hypothesized to correspond with reduced ANS and central nervous system activity. Evidence from ‘pure presence’ states suggests widespread reductions in brain electrical activity, contrasting with findings of increased gamma activity in earlier meditation phases, indicating distinct high-content vs. minimal perceptual content states (Boly et al., 2024).

Advanced meditation has been reported to induce a reduction in autonomic engagement, marked by low HRV despite unchanged or reduced heart rate (McCraty et al., 2009). Some practitioners can reach a so-called ‘breathless state’, characterized by marked reductions in both breathing rate and amplitude (Devarshi, 2010; Sparby, 2019). However, research on autonomic changes across meditation depth remains limited.

While deep meditation is often linked to ANS suppression, some techniques actively engage cognitive and emotional processing, influencing autonomic dynamics. Practices like Loving-kindness and Tonglen meditation, which involve high emotional absorption, may sustain HRV amplitude rather than suppressing autonomic function (Andreu et al., 2025; Kok et al., 2013). In this context, the extent of ANS involvement appears to depend on the form of embodiment cultivated by the practice, that is, the extent to which the practice engages body awareness. Some techniques emphasize bodily sensations and emotional engagement, thereby maintaining autonomic responsiveness, whereas other contemplative practices, particularly those emphasizing non-dual awareness and deconstructive inquiry, foster a dissolution of self and body awareness, which may attenuate autonomic signals. In these states, ordinary distinctions between self and body may fade, contrasting with practices that heighten embodied awareness (e.g., Loving-kindness or Tonglen). This distinction may help clarify how different contemplative techniques engage the ANS: practices emphasizing embodiment may sustain HRV amplitude through emotional absorption, whereas practices fostering disembodiment may attenuate autonomic responsiveness.

A model emerges in which STEs may unfold along a continuum of information-processing complexity (i.e high to low perceptual content states described above). On the high information processing complexity end of the spectrum, STEs are characterized by expanded sensory awareness associated at the neural level with increased gamma-band activity (Boly et al., 2024) and, as our results suggest, a potential increase in sympathovagal engagement. These features support a broadening of the sensory field, often accompanied by a felt sense of unity with the environment, enhanced relationality, and positive affect. According to the Jhanas progression framework, such high-content, metabolically demanding states may serve as gateways to deeper meditative absorptions, marked by progressive sensory withdrawal, dissolution of self-referential processing, and reduced physiological activity. Numadelic experiences are designed to facilitate both forms of STEs, integrating immersive, sensory-rich phases alongside progressive sensory reduction. However, achieving object-free pure awareness may require deeper states of absorption and nonegoic self-awareness (Metzinger, 2024), making deeper STEs less accessible to non-meditators, and likely explaining the relation between STEs and higher autonomic activity observed in our dataset. Future research should investigate the relation between HRV amplitude and STEs in advanced meditators, and the possibility that deeper forms of STEs may be associated with reduced rather than increased engagement.

### Limitations

#### Within-subject experience sampling

Although HRV amplitude appears to be a promising marker for STE, its ability to track moment-to-moment fluctuations within individuals remains uncertain. Given that our design included only two chapters during which ANS activity could reliably reflect subjective experience, free from external confounds such as body motion or non-natural breathing, it was not possible to assess whether HRV amplitude dynamically varied with STE ratings over time. It is worth noting that although some measures were adopted to minimize such confounds, motion and breathing practices are inherent to the positive impact of numadelics, by providing embodied practices which enhance presence and connection between participants. Future research efforts for physiological monitoring should therefore not aim to reduce these factors, but instead, incorporate structured rest periods between embodiment practices. This approach would help maximize the number of data points per participant, allowing for more robust assessments of ANS-STE interactions and enabling a deeper understanding of how physiological markers evolve throughout immersive experiences.

#### Mystical experience questionnaire

The MEQ-30 was used across both datasets in this study to assess the depth of participants’ STEs following either numadelic or psychedelic sessions. Notably, MEQ-30 ratings correlated with HRV amplitude in the psychedelic dataset, but not in the numadelic one. Widely considered the most commonly used questionnaire for assessing mystical or spiritual experiences, the MEQ-30 is derived from the earlier MEQ-43, itself grounded in Walter Stace’s seminal framework on the core features of spiritual awakening (Stace, 1960). However, the MEQ-30 was developed and psychometrically validated almost exclusively within psychedelic research contexts (Barrett et al., 2015). A recent study (Formoso, 2023) attempted to validate the MEQ-30 in a sample reporting non-psychedelic mystical experiences. The findings indicated that the scale’s factor structure did not adequately fit this population, suggesting potential phenomenological differences between psychedelic and non-psychedelic STEs.

Another important consideration is temporal specificity. In the numadelic dataset, our analysis focused on distinct experiential chapters, each accompanied by subjective ratings of ego reduction. In contrast, the MEQ-30 captured participants’ reflections on the overall experience, rather than its discrete phases, possibly introducing some noise in the relation between physiological measure and questionnaire scores. One might ask why a similar issue did not affect the observed relationship between HRV amplitude and STE ratings in the DMT dataset. In this case, we averaged HRV amplitude over a period following the acute sympathetic surge, thereby focusing the analysis on a post-‘ascent’ phase that phenomenologically corresponds to more stable emotional and metacognitive content (Timmermann et al., 2019).

#### Comparison with traditional meditation

The control meditation used in this study was not an established traditional practice, but rather a self-constructed adaptation of the numadelic experience, designed as an audio-guided meditation where participants internally visualized the scenes seen by the VR group. While it may be tempting to infer from the results that VR enhances meditative depth relative to non-VR meditation, it is important to acknowledge the possibility that a more authentic meditation practice may have been more effective at eliciting some degree of STE (NB: A thorough exploration of differences between numadelic VR and the control meditation will be addressed in a separate article). Given this limitation, our intent is not to claim that numadelic experiences surpass traditional meditatio§n in their ability to induce STEs. Instead, future research should reverse the approach, developing a VR experience based on an existing meditative tradition rather than constructing a meditation adapted from numadelic elements. This shift would allow for a more robust evaluation of whether numadelic meditation can function as a catalyst for contemplative practice—potentially enabling novices to access states of consciousness that would otherwise require years of dedicated practice to achieve.

## Conclusion

We introduced a novel measure of autonomic activity—HRV amplitude—that effectively captures inter-individual differences in the depth of STEs during high-absorption states, whether elicited through numadelic VR or psychedelic modalities. As a practical physiological marker, this measure opens new avenues for tracking transformative experiences as they occur.

While research into numadelic experiences remains in its early stages, empirical insights and participant feedback suggest a growing potential for these immersive journeys to evoke profound psychological shifts, offering a non-drug-based path for achieving states usually reserved for advanced contemplative practitioners. Understanding how best to guide individuals into NOSC—through gradual immersion that facilitates the softening of ego boundaries—will be essential. In this context, tools that can objectively reflect the unfolding of such states hold great promise, both for experience design and for biofeedback-enhanced therapeutic interventions. Looking ahead, further research may refine this measure and explore its integration into scalable contemplative and clinical practices, potentially making self-transcendence more accessible in secular contexts.

## Acknowledgements

This study was supported by the Imagine project obtained by MW funded by the Generalitat Valenciana (CISEJI/2022/46). VB was supported by a grant from the Tiny Blue Dot foundation awarded to DRG.

## Data availability

ECG data and audio file of the audio of the pre-recorded meditation ‘Ripple’ in English are available at https://doi.org/10.5281/zenodo.17061627.

Code for data-preprocessing, final dataset for statistical analyses (with questionnaires and subjective ratings as well as physiological measures of interest) and VR experience chapters video snapshots are available at https://github.com/vbonnelle/Ripple_Study_Bonnelle2025.

## Notes

https://doi.org/10.5281/zenodo.17061627

### Competing Interest Statement

VB and JW are founding board member, and JLH is co-founder and executive director at Numadelic Labs, a 501(c)(3) non-profit organization dedicated to initiate and conduct research involving numadelic technologies. JLH is CSO at aNUMA, a for-profit company providing access to numadelic VR experiences as part of its intervention for people with life threatening illnesses. DRG is a co-founder of aNUma, Inc. JW, MW, CA and AC are aNUMA facilitators and Contemplative practice trainers.

### Summary of Updates

This revised manuscript incorporates reviewers' suggestions. Some figures have been modified and the text has been edited at multiple loacations.

