## Supplementary Information for "Autonomic Indicators of Self-Transcendence: Insights from the Numadelic VR Paradigm"

**Table S1: Subjective rating of the experience**

| Dimension | English translation |
| --- | --- |
| Engagement | On a scale from 0 to 10, how would you rate your level of engagement (how involved you were during this sequence)? **Min = 0:** I found it difficult to participate **Max = 10:** I was completely absorbed in the experience |
| Valence | On a scale from -10 to 10, how would you rate the valence of your dominant emotion during this sequence? **Min = -10:** Unpleasant emotion(s) **Max = 10:** Pleasant/enjoyable emotion(s) |
| Intensity | On a scale from -10 to 10, how would you rate the intensity of your dominant emotion during this sequence? **Min = -10:** Low intensity **Max = 10:** High intensity |
| Boundaries | To what extent did you feel that the boundaries between your body and the external world were altered? **Min = 0:** The boundaries of my body did not change **Max = 10:** The boundaries of my body felt more diffuse and/or expanded beyond usual |
| Connection | Use the slider to describe the degree to which you felt connected with your session partner. **Min = -10:** Much less than I would feel under normal circumstances **Max = 10:** Much more than I would feel under normal circumstances |
| Ego reduction | To what extent did you feel that your sense of ego became smaller? **Min = 0:** My ego perception did not change **Max = 10:** My ego perception felt smaller than usual |

**Table S2: Descriptive statistics of autonomic measures**

|  | **Baseline** | **Chapter 4** | **Chapter 7** | **Post-session** |
| --- | --- | --- | --- | --- |
| **Means ± SD** |  |  |  |  |
| HR (bpm) | 75.8±11.8 | 76.7±11.6 | 74.8±11.7 | 73.3±10.9 |
| HRV amp (s) | 0.09±0.04 | 0.07± 0.02^N*^ | 0.10±0.06 | 0.09±0.05 |
| RMSSD (ms) | 50.9±13.7 | 51.1±11.7 ^N^ | 52.3±14.9 | 53.3±13.8 |
| PAI (a.u.) | 56.1±10.1 | 59.8±9.4 ^N^ | 65.6±10.3 | 57.5±8.9 |
| SAI (a.u.) | 51.3±17.7 | 49.5±16.5 | 47.1±16.5 | 48.7±16.9 |
| Resp amp (a.u.) | 0.06±0.03 | 0.10±0.18 | 0.11±0.17 | 0.07±0.05 |
| Resp rate (bpm) | 12.4±4.1 | 11.4±4.9 | 9.7±4.8 | 12.1±4.4 |

^N^ indicates a normal distribution (Shapiro test, p<0.05), * indicates significant difference between chapters 4 and 7 (Mann-Whitney test, p<0.05). SD=Standard deviation; a.u.=arbitrary units

**Table S3: Descriptive statistics of the measures of psycho-emotional state at baseline and post session**

|  | **VR** | | **Control** | |
| --- | --- | --- | --- | --- |
|  | **Baseline** | **Post-session** | **Baseline** | **Post-session** |
| **Means ± SD** |  |  |  |  |
| Positive affect | 10.5±2.7 | 13.8±2.2 | 9.3±2.7 | 12.5±2.0 |
| Negative affect | 4.4±2.7 | 2.9±2.4 | 5.4±3.1 | 3.8±2.9 |
| Relationality* | 6.7±1.8 | 9.5±2.1 | 7.1±1.6 | 8.7±1.8 |

*Indicates a significant group x time interaction (ANIVA). SD=Standard deviation
